# The SMC Hinge is a Selective Gate for Obstacle Bypass

**DOI:** 10.1101/2025.03.17.643644

**Authors:** Hon Wing Liu, Florian Roisné-Hamelin, Michael Taschner, James Collier, Madhusudhan Srinivasan, Stephan Gruber

## Abstract

DNA loop-extruding SMC complexes play vital roles in genome maintenance and DNA immunity. However, the ability of these ring-shaped DNA motors to navigate large DNA-bound obstacles, such as DNA/RNA polymerases and ribosomes in prokaryotes, has remained unclear. Here, we demonstrate that a bacterial SMC Wadjet complex can efficiently bypass obstacles larger than the SMC coiled coil lumen if they are linked to the extruded DNA by single-stranded DNA (ssDNA). This process is mediated by the selective entrapment of the ssDNA linker within the SMC hinge channel, which acts as an obligate gate—permitting ssDNA passage while excluding double-stranded DNA (dsDNA). By threading the linker through the hinge gate as dsDNA is translocated through the SMC motor, the obstacle can follow a trajectory distinct from its associated dsDNA, remaining outside the SMC ring. This mechanism enables continuous dsDNA entrapment while successfully bypassing obstacles. We show that hinges of diverse SMC complexes including eukaryotic condensin and Smc5/6 also selectively accommodate ssDNA, suggesting that hinge bypass is a conserved feature across SMC proteins. We demonstrate that while the hinge domain is essential for obstacle bypass, it is dispensable for loop extrusion, showing that these functions are independent, provided by separate SMC modules. These findings uncover the long-sought function of the SMC hinge toroid and provide the first mechanistic explanation for how DNA-entrapping SMC complexes can extrude loops on chromosomal DNA densely populated with obstacles. Integrating hinge bypass into the DNA loop extrusion model, we propose that the hinge may also facilitate the passage of RNA and peptide, enabling bypass of transcription-associated ribonucleoprotein complexes and other SMC complexes.

## Introduction

Chromosomes fold into compact yet dynamic 3D structures through DNA loop extrusion (LE), driven by conserved ATP-powered structural maintenance of chromosomes (SMC) motors. LE is the enzymatic activity that underpins diverse genome organizational features across life, from bacterial chromosome arm alignment to eukaryotic interphase chromosome domain formation and mitotic chromosome condensation. LE brings distant DNA segments into proximity to regulate transcription, repair, and recombination ^1–3^. SMC proteins harbor an extended coiled coil, with an ABC ATPase head domain at one end and a hinge domain at the other. They dimerize by forming a toroid-shaped hinge and bind a kleisin subunit at the heads, forming an elongated, ring-like structure ^4–6^. The complex is further elaborated by either a KITE dimer (in bacterial SMC complexes and in Smc5/6) or two HAWK subunits (in cohesin and condensin) that bind to the kleisin subunit ^7,8^. Purified SMC complexes can entrap and extrude DNA *in vitro* ^9–15^. While several models for the mechanism of LE have been put forward ^16–21^, the exact molecular processes of LE remains unclear.

Wadjet, a family of SMC protein complexes, are bacterial immune system components that use LE to selectively recognize and restrict invading plasmids by DNA cleavage ^22–25^ while sparing the larger chromosomal DNA (Extended Data Fig. 1A). Our previous work suggested that DNA is topologically entrapped inside Wadjet SMC rings during LE ^25^. If SMC-mediated LE serves as a universal mechanism for genome organization, it implies that the motors driving LE must navigate various obstacles on DNA, including nucleosomes, DNA replication and transcription machineries. In prokaryotes, ribosomes associated with transcription-translation complexes would also pose significant barriers to LE, further challenging the process. Although LE of “chromatinized” DNA has not been demonstrated *in vitro*, SMC motors have been observed to bypass individual nucleosomes. Strikingly, single-molecule studies show they can bypass other SMC complexes, forming interlocking “Z-loops,” and can even overcome artificial obstacles much bigger than themselves ^11,26,27^. Interestingly, the apparently efficient incorporation of large obstacles into DNA loops formed by permanently circularized SMC rings ^11,27^ led to the proposal that DNA loops might be held outside the SMC ring during LE ^19^. Hence, the need for and existence of bypassing remain debated, its mechanisms are unknown, and the specific components of SMC complexes responsible have yet to be identified. Here, we demonstrate efficient bypassing of large obstacles by an SMC complex and show that this process depends on access to ssDNA by the SMC hinge. We elucidate the SMC hinge as a selective gate, allowing passage of ssDNA but excluding dsDNA, which is a conserved feature among diverse SMCs. We find that LE and obstacle bypass activities by SMC complexes are independent of each other and are contributed by different parts of the SMC ring.

### A ssDNA requirement for efficient obstacle bypass by Wadjet-I

Wadjet (JetABCD) SMC complexes restrict plasmids through DNA cleavage in diverse bacteria and archaea. They exist in three types (I, II, and III) with differences in operon organization and domain composition ^24,28^. Cleavage is DNA sequence non-specific and involves a dimeric JetABC motor actively extruding the plasmid, followed by activation of the JetD nuclease subunit once extrusion is complete (Extended Data Fig. 1A) ^13,22,23,25^. We exploit this extrusion-cleavage activity to study roadblock bypassing, with DNA molecules cleaved at the roadblock position marking extrusion-cleavage events that occurred without bypassing events (Fig. 1A). Wadjet-I (from *Escherichia coli* strain GF4-3, ‘ecJET’) cleaved DNA circles anchored to micrometer-scale Dynabead obstacles via streptavidin-biotin linkage mostly at the obstacle anchor and only occasionally away from the anchoring point ^25^. This observation raised the question of whether the enzyme’s inefficiency in bypassing the obstacle was a consequence of constraints imposed by the synthetic DNA-obstacle linkage itself ^25^.

**Fig. 1.**
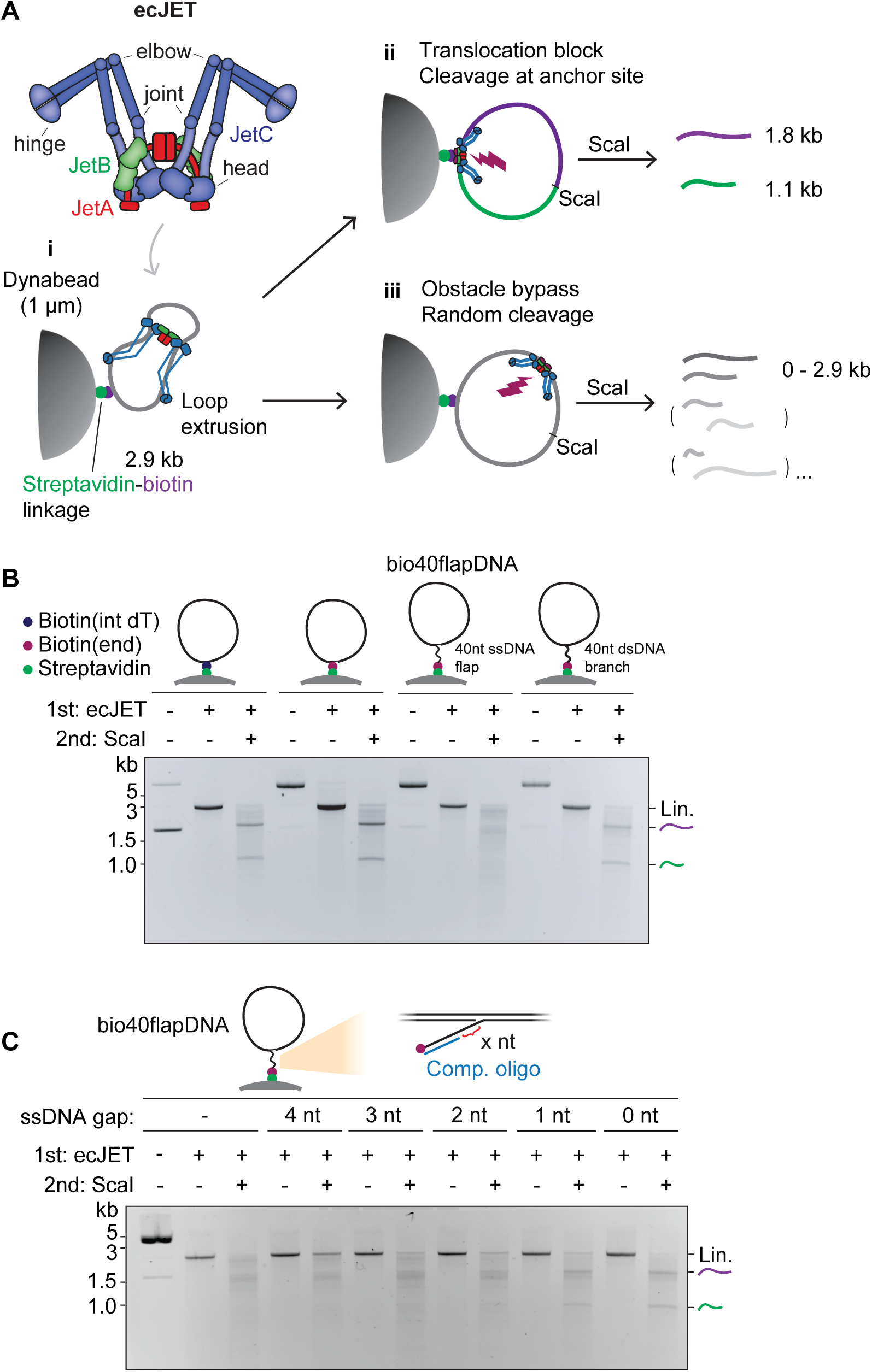
- Requirements for obstacle bypassing by Wadjet-I. A) Schematic depiction of Wadjet-I from *E. coli* strain GF4-3 (ecJET). JetC (blue) forms the SMC dimer, JetB (green) is the KITE dimer, JetA (red) is the kleisin subunit, altogether forming a dimer-of-motors configuration. The JetC hinge domain and elbow are highlighted. i) Depiction of cleavage and roadblock bypass assay by Wadjet, on 2.9 kb DNA circles containing a Dynabead obstacle (attached via biotin-streptavidin linkage). Wadjet loads onto DNA, then extrudes a loop, followed by one of two possible outcomes: ii) Wadjet stalls at the obstacle, triggering cleavage at the roadblock anchor point. Post-treatment by ScaI single cutter results in two fragments 1.8 kb (purple) and 1.1 kb (green) in size. iii) One motor unit of Wadjet bypassing the obstacle, resulting in cleavage at random position. Post-treatment by ScaI results in fragments of variable size. See Extended Data Fig. 1A for the Wadjet plasmid restriction mechanism on naked DNA. B) Wadjet bypasses a large obstacle (Dynabead) attached onto DNA by a ssDNA linker. Agarose gel showing the cleavage activity of ecJET on DNA circles containing biotin moieties at various indicated positions, attached onto streptavidin-coated Dynabeads. Two bands migrating at 1.8 and 1.1 kb indicate DNA cleavage at the roadblock anchor while a smear indicates random cleavage after obstacle bypass. Note that untreated DNA circles containing 5’-positioned biotins migrate slower, as they contain a DNA nick and thus are open covalent circles (occDNA), circles with internally positioned biotin are closed (cccDNA) and thus migrate faster. See Extended Data Fig. 1B for details on biotinylated DNA substrate generation. C) Agarose gel showing the cleavage activity of ecJET on bio40flapDNA attached onto Dynabeads with a partially complementary oligo (in blue) annealing at the ssDNA flap, leaving behind a defined length of ssDNA.

To investigate this further, we systematically modified the positioning of the biotin moiety on the 2.9 kb DNA circle (Fig. 1B, Extended Data Fig. 1B). ecJET tended to cleave at the Dynabead obstacle attachment point on DNA circles containing biotin positioned at the 5’ end of a nick, as previously seen with internal biotin ^25^. Strikingly, the cleavage site distribution became random if forty ssDNA nucleotides were placed between the plasmid DNA and the 5’ end biotin (hereafter abbreviated as bio40flapDNA) (Extended Data Fig. 1B), indicating that the Dynabead surprisingly posed little or no barrier for ecJET if attached via ssDNA. Given that extrusion of this short DNA substrate is expected to occur within seconds, this finding implies that bypassing is not only efficient but also fast. Notably, bio40flapDNA circles were as stably anchored on Dynabeads as in the other scenarios (Extended Data Fig. 2A), making it unlikely that the random cleavage pattern was due to DNA dissociation from the beads. Remarkably, converting the ssDNA flap to a dsDNA branch by annealing a complementary oligonucleotide reverted the cleavage pattern by ecJET to an anchor-site-specific one (Fig. 1B). This suggests that ecJET can indeed efficiently bypass DNA obstacles larger than its own dimensions—as long as the linkage to the DNA includes a certain extent of single-stranded DNA. By annealing shorter oligonucleotides (thus resulting in dsDNA-ssDNA hybrid flaps, Extended Data Fig. 1B), we found that a stretch of ten ssDNA nucleotides at the biotin end was sufficient to create a permeable barrier. Intriguingly, even two ssDNA nucleotides at the plasmid end enabled efficient bypass (Fig. 1C, Extended Data Fig. 2B). This suggests that bypass can occur both immediately adjacent to the plasmid DNA and at a distance from it, with the former configuration being more favorable for SMC bypass in our assay. Importantly, ssDNA stretches that were not part of the linker but were positioned adjacent to the bead anchor, such as a 10-nucleotide 3’ flap or a 12-nucleotide bubble near the biotin moiety, did not facilitate efficient bypass (Extended Data Fig. 2C). This highlights the critical role of ssDNA specifically within the linker region, rather than its mere presence near the anchor, in enabling ecJET to navigate around obstacles.

### Wadjet-I hinge domains facilitate DNA obstacle bypass

SMC proteins typically adopt a donut-shaped hinge structure featuring a central channel lined with positively charged residues (Fig. 2A, 3A, Extended Data Fig. 8). Despite the high conservation of these charged residues across species, their specific functional role remains poorly understood. Intriguingly, this feature is absent from some SMC proteins including a Wadjet-II (from *Neobacillus vireti* strain LMG 21834, ‘nvJET’) (Fig. 2B, Extended Data Fig. 8), which, like ecJET, is capable of cleaving naked circular DNA ^29^. Strikingly however, nvJET was totally bypass-incompetent, even with bio40flapDNA (Fig. 2B), resulting in DNA cleavage exclusively at the obstacle anchor position. This raises the interesting possibility that only hinges with a donut shape might be capable of facilitating obstacle bypass by temporarily accommodating ssDNA in their channel and acting as single-strand-specific gates. To determine whether hinge domains must detach from one another for obstacle bypass, we utilized a Cys-less ecJET (that lacks all endogenous cysteine residues; manuscript in preparation) and introduced cysteine pairs at ecJetC hinge interfaces. Specifically, we engineered two derivatives containing either K529C/R593C or Q508C/H576C, enabling closure of the hinge donut via chemical crosslinking with 1,4-butanediyl bismethanethiosulfonate (M4M) (Fig. 2A, Extended Data Fig. 3A) ^30^. Cys-less ecJET (WT) was proficient in plasmid cleavage and efficiently bypassed Dynabeads on bio40flapDNA (Fig. 2C). While hinge crosslinking in both cysteine pair variants did not hinder DNA cleavage, it completely abolished the enzyme’s ability to bypass Dynabeads on bio40flapDNA. Notably, random cleavage was restored when the crosslinked Wadjet proteins were treated with the reducing agent DTT, which reopened the hinge (Fig. 2C, Extended Data Fig. 3B). Hinge closure also prevented random cleavage on all other DNA substrates tested except for obstacle-free bio40flapDNA (Extended Data Fig. 3C-D). These findings unambiguously assign the hinge domain in Wadjet-I as a gate facilitating “hinge bypass” of ssDNA-anchored obstacles. Wadjets harboring closed hinges, whether naturally occurring or artificially crosslinked, also failed to efficiently cleave DNA circles with two bio40flaps for bead attachment (Fig. 2D, Extended Data Fig. 3E). This indicates that hinge bypass is important for complete DNA extrusion and subsequent cleavage to occur, particularly in this double-obstacle scenario. Together, these findings highlight the critical role of hinge domain in enabling loop extruding Wadjet-I to navigate around obstacles.

**Fig. 2.**
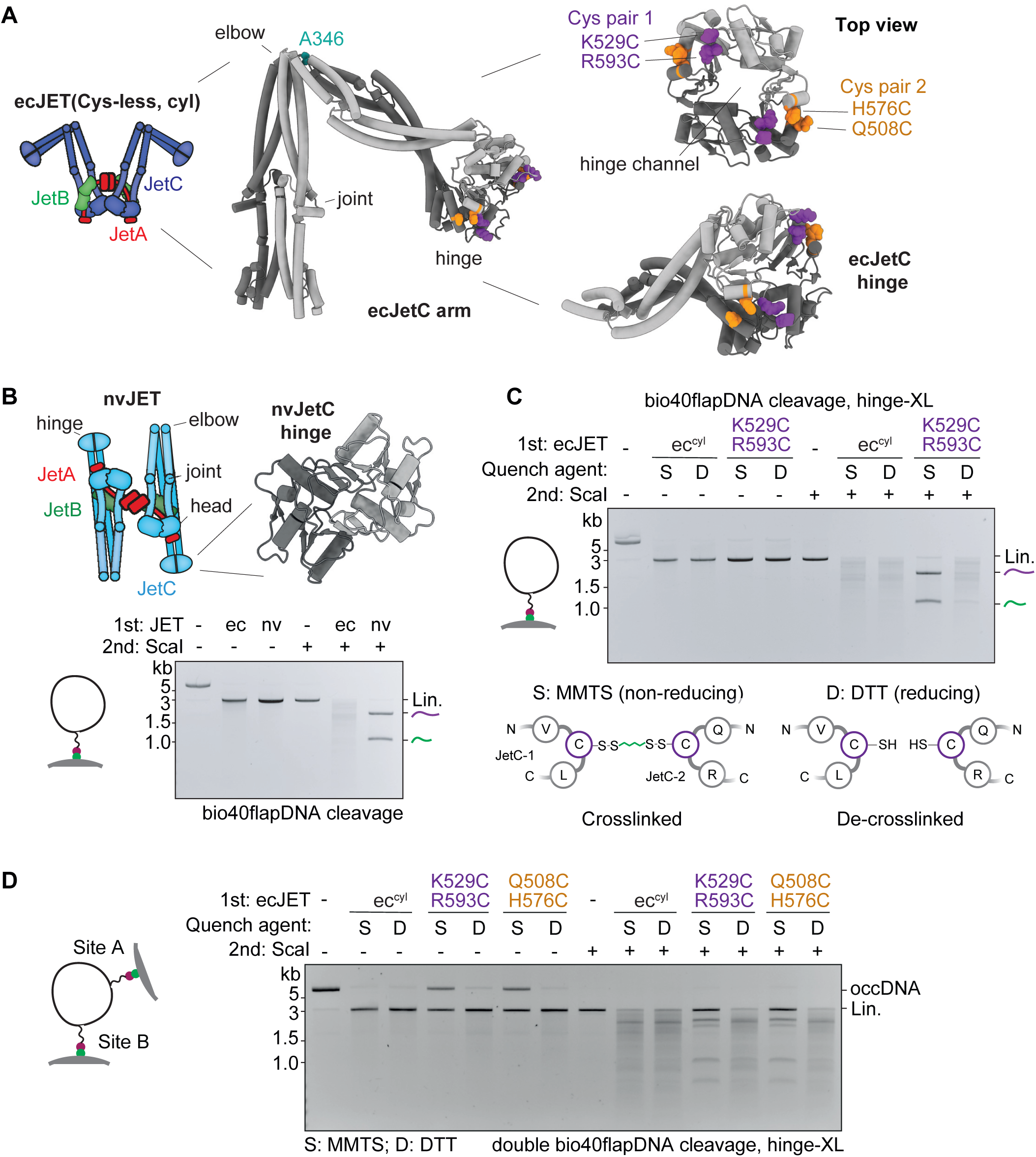
- The hinge domain of Wadjet-I facilitates obstacle bypassing. A) Left: Structural model of the ecJetC arm, with the A346 residue at the elbow position used for crosslinking highlighted in teal colors. Right: zoomed in views of the ecJetC hinge, with the two cysteine pairs for crosslinking (pair 1: R529, R593 in purple colors; pair 2: Q508, H576 in orange colors) highlighted. The hinge channel is recognizable in top view. B) Schematic of Wadjet-II nvJET (*N. vireti* strain LMG 21834) and a close-up model of its channel-less hinge dimer ^29^. Bottom: Agarose gel depicting the cleavage/bypass activity on bio40flapDNA attached onto Dynabeads by the indicated Wadjets (ec, ecJET; nv, nvJET). C) Top: Agarose gel depicting the cleavage/bypass activity on bio40flapDNA attached onto Dynabeads by the indicated ecJET variants, after M4M-induced cysteine crosslinking and subsequent quenching by the non-reducing *S*-Methyl methanethiosulfonate (MMTS) quencher (preserving the crosslinks) or reducing dithiothreitol (DTT) (disrupting the crosslinks), depicted in the bottom panel (see Methods for details). “ec^cyl^” denotes Cys-less ecJET (manuscript in preparation). The hinge crosslinking mutant contains two additional cysteine substitutions at K529 and R593 (pair 1). See Extended Data Fig. 3A for a measure of the degree of crosslinking. See panel D and Extended Data Fig. 3B for experiments with Cys-less ecJET harboring another hinge crosslinking cysteine pair at positions Q508 and H576 (pair 2). D) Agarose gel showing the cleavage/bypass activity of the indicated ecJET variants on DNA circles with two bio40flap-Dynabead modifications. ecJET containing chemically crosslinked hinges cleaved these substrates less efficiently as they are not able to bypass the two obstacles to reach a cleavage-competent state. Residual cleavage events were likely due to a population of DNA circles that have only attached onto a streptavidin Dynabead at one of the two sites, which do allow cleavage by hinge-crosslinked ecJET at the Dynabead anchor, as evidenced by the resultant four defined smaller DNA fragments after ScaI post-treatment. We also note that DNA cleavage site distribution by uncrosslinked ecJET was unperturbed (kept random) by the presence of two obstacles. See Extended Data Fig. 3E for the experiment with Wadjet containing naturally closed hinges. occDNA - open covalent circles.

### ssDNA entrapment by SMC hinge domains

We further characterized the SMC hinge, by first investigating whether the highly conserved positive charges that line the ecJetC hinge channel are important for hinge bypass, presumably through charge interactions with the ssDNA linker. To this end, we generated ecJET complexes with triple alanine (3A) and aspartate (3D) substitutions of ecJetC hinge channel residues R524, K591, K601 (Fig. 3B). The 3D mutant was completely defective in Dynabead bypassing but also displayed a defect in DNA cleavage, whereas the 3A mutant exhibited a significant reduction in bypassing activity whilst retaining normal DNA cleavage efficiency (Fig. 3B), together strongly supporting the notion that the channel charges facilitate hinge bypass by capturing ssDNA. We then tested the ability of the 3A mutant to restrict a 4 kb test plasmid *in vivo* and found that the residual bypass activity of the 3A mutant was sufficient for restriction of this substrate (Extended Data Fig. 4A). We speculate that the positive charges play a more critical role in restricting larger, more complex natural Wadjet substrates.

**Fig. 3.**
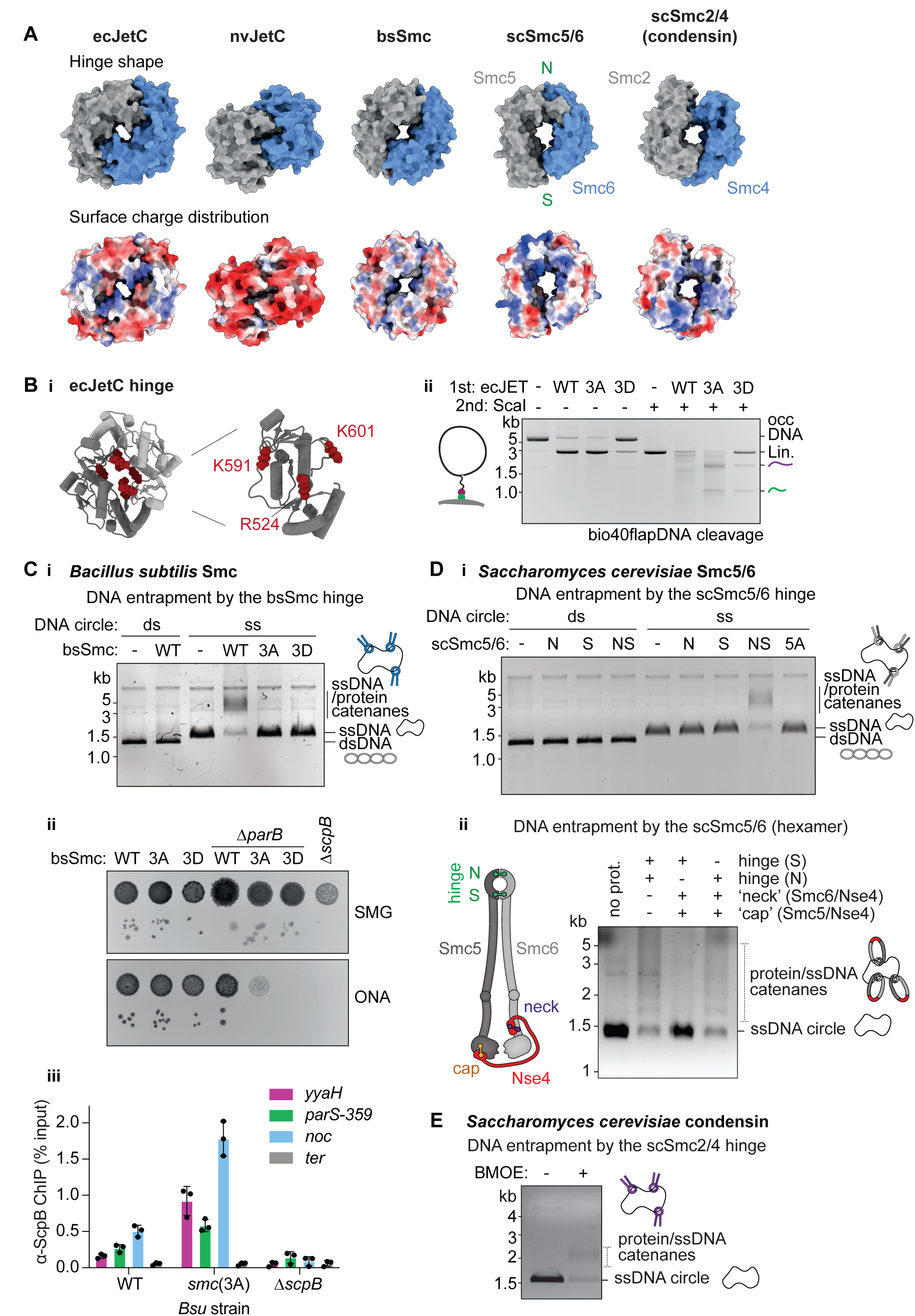
- ssDNA entrapment by SMC hinges. A) Top: Top view structures of selected hinge dimers used in this study. ecJetC, nvJetC and bsSmc - AlphaFold predictions. scSmc5/6, PDB: 7QCD ^57^; scSmc2/4, PDB: 6YVU^58^. Note the absence of a cavity/channel in the nvJET hinge. The “North/N” and “South/S” interfaces used in subsequent crosslinking experiments are marked for the scSmc5/6 hinge. Bottom: Surface charge distribution (red indicating more negatively charged; blue indicating more positively charged) of the same hinges. Note that the nvJET hinge also lacks any discernible positive charges at the dimer interface. See Extended Data Fig. 8 for depictions of other notable SMC hinges. B) i) Structural model of ecJET hinge depicting (in red colors) positively charged channel residues used for mutagenesis. ii) Agarose gel depicting the cleavage activity on bio40flapDNA by the indicated ecJET variants. 3A: ecJetC(R524A, K591A, K601A); 3D: ecJetC(R524D, K591D, K601D). See Extended Data Fig. 4 for the *in vivo* plasmid restriction phenotypes of these mutants. occDNA - open covalent circles. C) i) DNA entrapment by the bsSmc hinge. Denaturing agarose gel depicting 2.3 kb circular ss/dsDNA entrapment by the indicated bsSmc hinges (residues 400-776). All hinges harbor C437S to mitigate non-specific crosslinking as well as engineered cysteines R558C and N634C for crosslinking by BMOE. 3A/3D contain additional alanine or aspartate substitutions (respectively) at R583, R643, R645 positions of the hinge channel. ii) Colony formation assay on minimal (SMG) or rich (ONA) media of the indicated *B. subtilis* ssDNA entrapment deficient hinge mutant strains. 3A: bsSmc(R583A, R643A, R645A), 3D: bsSmc(R583D, R643D, R645D). Failure to grow on rich medium is caused by a defect in chromosome segregation and a lethal accumulation of interlinked sister chromosomes in the absence of sufficient SMC activity ^59^. iii) Chromatin immunoprecipitation coupled to real-time quantitative PCR (ChIP-qPCR) using a-ScpB serum with the indicated *B. subtilis* strains and primer pairs for four different loci. Individual data points are shown as dots and error bars depict standard deviation from three independent repeats. 3A: Mutant containing bsSmc(R583A, R643A, R645A). D) i) DNA entrapment by the scSmc5/6 hinge. Denaturing agarose gel depicting 2.3 kb circular ss/dsDNA entrapment by the indicated Smc5/6 hinges (Smc5 residues 305-805, Smc6 residues 407-808), crosslinked by BMOE. N: “North” interface containing Smc5(V638C), Smc6(N572C); S: “South” interface containing Smc5(N526C), Smc6(N643C). NS: Mutant containing N and S mutations. 5A: NS mutant containing hinge channel mutations Smc5(R537A, K612A, K641A), Smc6(R555A, R676A). ii) DNA entrapment by the scSmc5/6 hexamer. Left: Schematic of the tripartite Smc5/6 ring subunits with the crosslinked interfaces marked. Right: Denaturing agarose gel depicting 2.3 kb circular ssDNA entrapment by the scSmc5/6 hexamer after crosslinking the hinge (N/S interface) in combinations with crosslinking the neck and cap interfaces. See Extended Data Fig. 6 for a detailed explanation of the likely ssDNA trajectory into the Smc5/6 ring inferred from this data. E) Agarose gel showing 2.3 kb circular ssDNA entrapment by the condensin scSmc2/4 hinge (Smc2 residues 443-740, Smc4 residues 598-923), containing cysteine residues Smc2(S560C, K639C), Smc4(V721C, M821C) for closure by BMOE crosslinking.

Building on this, we sought to demonstrate ssDNA capture within the SMC hinge toroid and asked whether it is a universal mechanism shared by SMC proteins from diverse species, despite their varied biological functions. Interestingly, predictions from AlphaFold3 imply a role for these residues in stabilising hinge-ssDNA interaction (Extended Data Fig. 4B). We purified isolated hinge domains from *B. subtilis* Smc (bsSmc), *S. cerevisiae* Smc5/6 (scSmc5/6), and condensin (scSmc2/4) as well as ecJET ^10,31^. Notably, the asymmetric scSmc2/4 and scSmc5/6 hinge have distinct “North” and “South” interfaces, while in the homodimeric bacterial hinges, the two interfaces have identical sequences (Fig. 3A). We incubated the purified hinge domains with either plasmid DNA or phage-derived 2.3 kb ssDNA circles, followed by crosslinking using bismaleimidoethane (BMOE), and analyzed the reaction products by protein-denaturing agarose gel electrophoresis. We found that crosslinked bsSmc and scSmc5/6 hinges entrapped ssDNA but not dsDNA circles (Fig. 3C-D, Extended Data Fig. 5A-B). scSmc2/4 as well as the ecJetC hinge also displayed ssDNA entrapment in this assay (Fig. 3E, Extended Data Fig. 4B, 5D). Additionally, we observed that mutating positively charged residues in the hinge channel of both bsSmc and scSmc5/6 hinges abrogated ssDNA entrapment (Fig. 3C-D). These findings support the notion that capture of ssDNA inside the hinge toroid is a conserved property of all SMC complexes and that the positively charged residues lining the channel are essential for this.

We next asked how ssDNA enters the hinge toroid, and if ssDNA preferentially traverses through one of the two hinge interfaces. This is difficult to test in SMC complexes containing symmetric hinge domains. We therefore exploited the asymmetry of the scSmc5/6 hinge by performing ssDNA entrapment experiments in the context of the hexameric core complex ^10^. We created chemically closed compartments by individually pairing a hinge cysteine residue (North or South interface) with two cysteine pairs at the kleisin/SMC interface. Strikingly, we found that only the North combination led to efficient ssDNA entrapment (Fig. 3D, Extended Data Fig. 5E), suggesting that ssDNA likely uses the North interface as a primary gate for entering the hinge lumen (Extended Data Fig. 6). Whether this is a common feature in all asymmetric hinge containing SMCs and whether ssDNA linked with obstacles follow the same trajectory through the hinge remains to be established.

SMC complexes on the *B. subtilis* chromosome are thought to efficiently bypass transcription units and other SMC complexes ^32–34^. We examined the effects of mutating the bsSmc hinge channel. While strains with bsSmc hinge mutations (3A, 3D) showed no significant growth defects alone (Fig. 3C), they exhibited strong synthetic phenotypes when combined with a chromosome segregation mutation (Δ*parB*), indicating impaired segregation without hinge bypass. Chromatin immunoprecipitation (ChIP) of a bsSmc(R583A, R643A, R645A) strain revealed increased KITE subunit ScpB occupancy at origin-proximal loci (Fig. 3C), suggesting retarded translocation and accumulation of bsSmc near *parS* sites possibly due to lost hinge bypass activity, likely affecting obstacle navigation.

### The SMC periphery is required for obstacle bypass but dispensable for loop extrusion

We have now established the critical role of the SMC hinge in facilitating obstacle bypass during extrusion by ecJET. However, an important question remains: does the hinge domain also play a role in extruding obstacle-free DNA? To test this, we engineered three variants of ecJET, focusing on the JetC elbow, a folding point in the middle of the SMC coiled coil ^25,35^. First, we wondered whether substitution of the ecJetC elbow-to-hinge segment for the corresponding (and much larger) nvJetC segment would abolish LE and obstacle bypass. Such a chimera (‘chim’) was readily expressed and purified (Extended Data Fig. 7A). It retained efficient DNA cleavage (Fig. 4A), suggesting that LE activity remained intact despite the lack of a cognate elbow-hinge segment. However, it failed to bypass Dynabeads on bio40flapDNA, presumably due to the presence of the bypass-incompetent nvJET hinge (Fig. 4A). Next, we investigated whether the formation of an open elbow-to-hinge lumen is required for LE and obstacle bypass by introducing a crosslink at the JetC elbow (A346C) in Cys-less ecJET. Crosslinking at this site also blocked obstacle bypass (Fig. 4B) but had minimal impact on DNA cleavage. These findings reinforce the notion that the hinge and the hinge proximal coiled coil must remain openable for obstacle bypass. Crucially, they suggest that neither the hinge nor the associated coiled coil is necessary for LE.

**Fig. 4.**
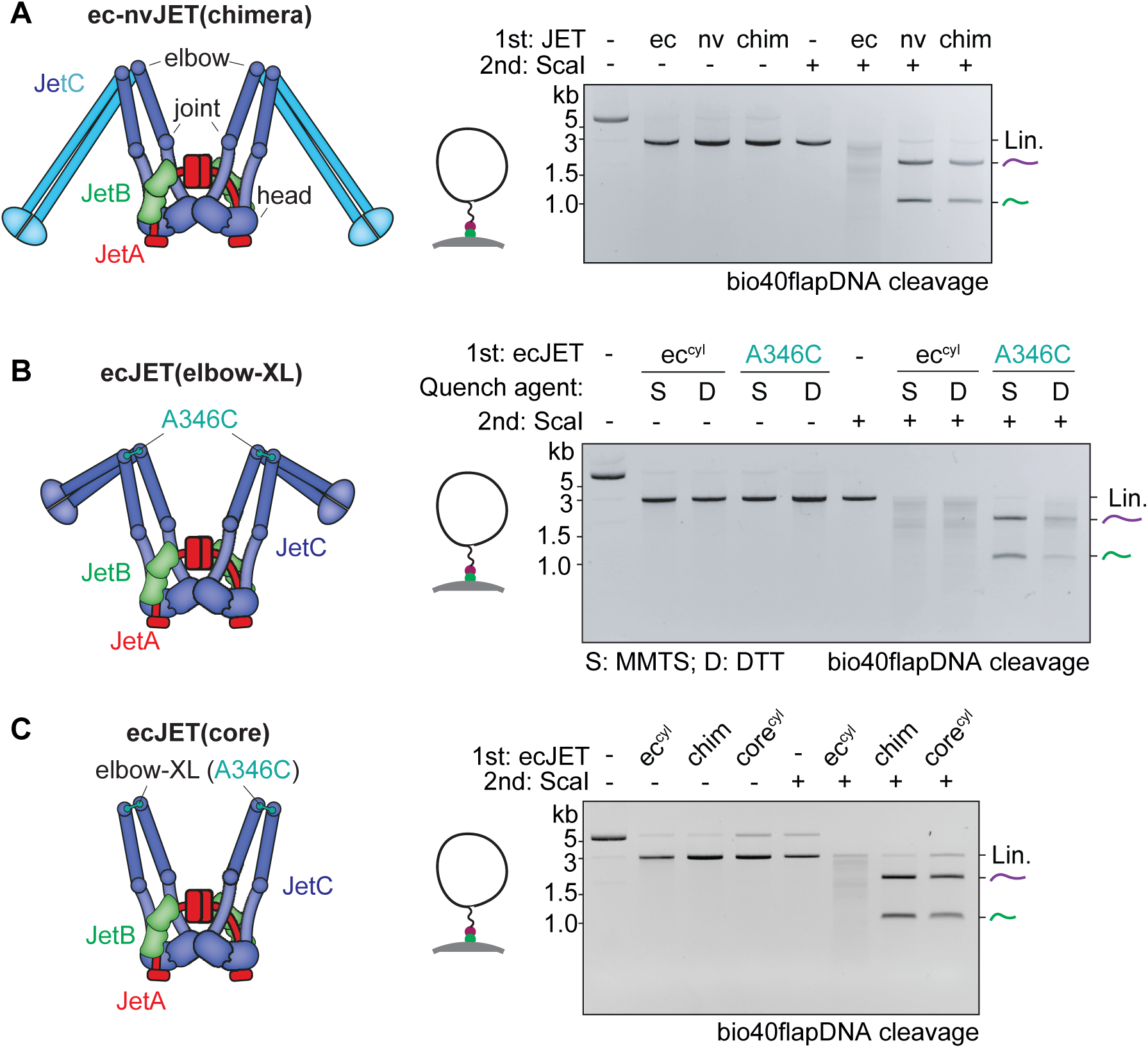
- The SMC periphery is required for obstacle bypassing but not DNA loop extrusion. A) Dynabead bypass experiment on bio40flapDNA using a chimera (‘chim’) harboring the ecJET core and an nvJET elbow-to-hinge segment. B) As with A but crosslinking the ecJET elbow position (A346C) in a Cys-less ecJET background (ec^cyl^). C) As with A but with a Cys-less ecJET(core) complex lacking the elbow-to-hinge segment (‘core’). The elbow-to-hinge segment is replaced by a nine-residue linker peptide (not shown).

To definitively rule out a requirement of the elbow-to-hinge segment for LE, we created a Cys-less ecJET core-only complex (‘core’) replacing the entire ecJetC elbow-to-hinge segment (aa 348-686) with a nine-residue linker peptide (GGGGSGGGG). To ensure stable dimerization of ecJetC(core) in the absence of a hinge, we included the elbow crosslink cysteine (A346C) to allow disulfide bond formation when purifying it under non-reducing conditions (Extended Data Fig. 7B). Remarkably, ecJET(core) retained the ability to cleave plasmid DNA but was completely unable to bypass obstacles (Fig. 4C). As expected, pre-treatment with DTT eliminated DNA cleavage in this scenario (rather than rescuing bypass), indicating that the SMC ring must be closed for LE to occur (Fig. 4C, Extended Data Fig. 7C). Collectively, these findings suggest that SMC complexes consist of a core module capable of extruding naked DNA and peripheral region (the elbow-to-hinge segment; ‘SMC periphery’) that independently facilitates navigation on complex DNA substrates by enabling obstacle bypass (Fig. 5A, B). This modular organization of SMC ring function, with distinct regions dedicated to different tasks, must be integrated into any comprehensive model of LE.

**Fig. 5.**
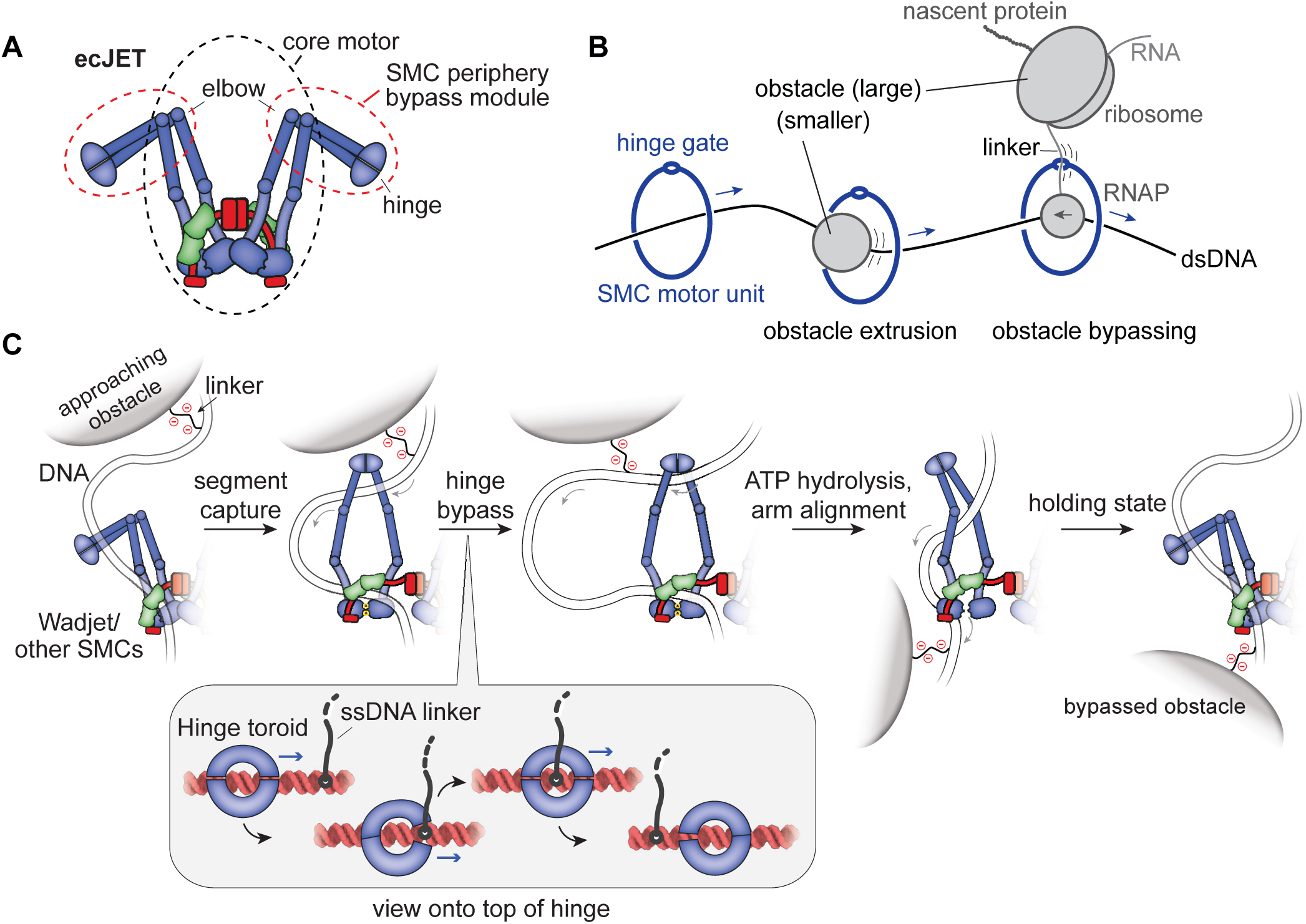
- A model for obstacle bypass by DNA loop-extruding SMC complexes. A) Distinct regions of the SMC complex responsible for loop extrusion (LE) and for obstacle bypass. The core for basic loop extrusion includes the head module and the head-to-elbow coiled coils. The periphery for obstacle bypass comprises the hinge and the adjacent elbow-to-hinge coiled coils. B) Two pathways for obstacle bypass: threading and hinge gating. C) The segment capture+ model builds on the DNA segment capture mechanism, which includes two alternating steps: the DNA holding state—with ATP-disengaged heads, closed coiled coil lumen and DNA held by the kleisin compartment—and the segment capture state—with ATP-engaged heads, open coiled coil lumen and a segment of DNA held by the SMC compartment and clamped at the heads by the kleisin subunit. ATP hydrolysis converts the segment capture state into the DNA holding state. The segment capture+ model incorporates an additional step for obstacle bypass in the segment capture state. The stepwise passage of the obstacle linker through the double-gated hinge is illustrated in the close-up view. For simplicity, only a single motor unit is shown.

## Discussion

Loop-extruding SMC complexes encounter various obstacles on their chromosomal translocation tracks. Our findings define two distinct pathways for overcoming roadblocks, (i) obstacle extrusion by threading through the SMC lumen and (ii) specific obstacle bypass via the hinge gate (Fig. 5B). Smaller obstacles are putatively readily overcome by threading through the SMC lumen, such as transcription factors and nucleosomes, while others are too large for threading, exceeding the lumen size, including transcription-translation complexes in bacteria and transcription-splicing complexes in eukaryotes. The identification of the hinge as bypass gate elucidates how SMC motors navigate these larger obstacles, effectively removing size limitations provided that the obstacles contain linkages that can be accommodated by the hinge (Fig. 5A, B). Because the hinge gate allows passage only of a single nucleic acid strand at a time, the SMC ring remains sealed for dsDNA, even on obstacle-laden DNA.

### The hinge bypass gate

We define functionally separable modules within SMC complexes: a core complex responsible for DNA loop extrusion and an SMC periphery comprising the hinge and adjacent coiled coils for obstacle bypass (Fig. 5A). Our findings link the characteristic donut-shaped structure of the hinge^5^ and its conservation to its role in selectively accommodating stretches of single-stranded nucleic acids and in passing them from one side of the ring to the other through a dual-gated mechanism (Fig. 5A-B). This provides a critical advantage for SMC complexes manoeuvring the crowded chromosomal environments, allowing them to get across challenging genomic regions effectively.

While our study revealed ssDNA linkages as efficient substrates for hinge bypass by ecJET, the observation that linkages in the form of carbon chains, lacking DNA (Fig. 1B, Extended Data Fig. 1B, 3C), still permit bypassing, albeit inefficiently, suggests that SMC hinges exhibit a degree of promiscuity. RNA molecules feature similar chemistry, size, and charge distribution to ssDNA. We suspect that in cells, transcription-based ribonucleoprotein complexes— particularly at highly transcribed genes—serve as the primary substrates for hinge bypass. Moreover, the SMC-SMC bypassing proposed for condensin based on single-molecule imaging ^26^ and inferred for bsSmc from its chromosomal dynamics ^32,34^ could be explained if the respective hinge gates also accommodate peptides that mimic ssDNA. Peptides suitable for hinge bypass may be present in the extended unstructured regions of kleisin subunits, particularly in cohesin and condensin. Notably, a motor unit from one LE complex bypassing a motor unit from a neighboring LE complex could explain the formation of interlocking ‘Z-loops’ ^26,32,34^. Supporting the concept of hinge-mediated SMC-SMC bypass, hinge-substituted bsSmc proteins exhibit defects in chromosome folding. These defects can be suppressed by reducing the number of *parS* loading sites on the chromosome to one, thereby minimizing the likelihood of SMC-SMC collisions ^32^.

### Two pathways for obstacle bypassing and implications for SMC evolution

Not all SMC complexes employ hinge bypass. Wadjet-II, which has lost the central channel in its hinge, is unable to bypass a ssDNA-linked obstacle in our assay. Interestingly, Wadjet-II, along with MukBEF (Extended Data Fig. 8) and Wadjet-III—both of which also lack a hinge channel—tend to have significantly longer coiled coils (with few exceptions in Wadjet-III). Similarly, Smc proteins from *Acetobacter* and related genera lack a hinge channel as predicted from AF3 (Extended Data Fig. 8) and have unusually long coiled coils (with an amino-terminal coiled coil a-helix of 511 and 512 aa in *Acetobacter pasteurianus* and *Gluconacetobacter diazotrophicus*, respectively, compared to 330 aa in bsSmc)^36^. The substantial enlargement of the SMC periphery (Fig. 5B) likely enables the bypassing of larger obstacles by threading them directly through an expanded SMC lumen. However, it is unlikely that lumen expansion—itself limited by finite protein sizes—fully compensates for the loss of hinge bypass. We speculate that anti-Wadjet-I counter defence mechanisms may have specifically targeted the hinge channel, which gave Wadjet-II and -III a selective advantage over other Wadjet systems in such environments.

In *Bacillus subtilis*, the two pathways of bsSmc obstacle bypass function alongside an obstacle avoidance mechanism mediated by the ParB protein, which loads bsSmc complexes onto the chromosome at *parS* sites near the replication origin. bsSmc loading at *parS* sites aligns their translocation with replication forks and most transcription units, minimizing the risk of head-on collisions ^32,33^. Consequently, *parB* deletion increases reliance on hinge bypass (Fig. 3C). The presence of three distinct strategies to mitigate roadblocking underscores their physiological significance and highlights the need to investigate this process also in more complex organisms.

### Additional functions of the hinge gate in cohesin and Smc5/6

Smc5/6 has been proposed to anchor onto ssDNA to facilitate DNA repair processes, for example stabilizing ssDNA/dsDNA junctions ^43,44^. *In vitro*, an interaction between cohesin and R loops slows down DNA translocation ^45^. These observations might be explained by prolonged ssDNA/RNA entrapment by the respective hinges. The donut-shaped cohesin hinge (Extended Data Fig. 8) is also implicated in the establishment of sister chromatid cohesion in yeast ^37–39^ potentially involving hinge bypass of ssDNA at replication forks ^40^. Supporting this, cohesin hinge channel mutants exhibit strong cohesion defects in yeast and humans ^41,42^. Accordingly, the hinge bypass gate may serve as a DNA entry gate in cohesin ^37^.

### Outlook

By characterizing an archetypical SMC complex proficient in LE, we have identified a key molecular determinant of obstacle bypass. While the findings remain agnostic to any proposed model of DNA loop extrusion, we note that the hinge bypass integrates seamlessly into the segment capture model ^16,17^ (Fig. 5C). Our data demonstrate that LE proceeds uninterrupted even when the SMC arms are unable to fold at an elbow. Furthermore, we show that the hinge and the proximal coiled coils, up to the elbow, while essential for obstacle bypass, are entirely dispensable for LE itself. These findings challenge other popular models that require repeated elbow folding and unfolding of SMC coiled coils for LE and call for a reassessment of such mechanisms. Our discovery that single-stranded nucleic acids are critical for obstacle bypass opens intriguing possibilities for establishing a molecular framework for further exploration of fundamental activity in eukaryotic cells: namely how cohesin can accommodate the replication fork to generate cohesion and how condensin is able to assemble mitotic chromosomes.

## Acknowledgments

We are grateful to Stéphane Marcand and members of the Gruber Lab for critical feedback on an earlier version of the manuscript and for stimulating discussions.

## Funding

This study was supported by the European Research Council (724482 to S.G.) and by the Wellcome Trust (226494/Z/22/Z to M.S.). H.W.L. and F.R.-H. were supported by EMBO Postdoctoral fellowships (ALTF 490-2021 and ATLF 302-2022).

### Author contributions

Formal analysis: H.W.L., F.R.-H., M.T.; Investigation: H.W.L., F.R.-H., M.T., J.C.; Methodology: H.W.L., F.R.-H., M.T., J.C.; Materials: core-, chim-, cysless-ecJET: F.R.-H.; condensin hinge: J.C.; other hinges: M.T., H.W.L., F.R.-H.; DNA substrates: H.W.L., M.T.; Supervision: S.G., M.S; Visualization: H.W.L., F.R.-H., M.T., S.G.; Writing – original draft: H.W.L., S.G.; Writing – review & editing: H.W.L., F.R.-H., M.T., J.C., M.S., S.G; Funding acquisition: S.G., H.W.L., F.R.-H., M.S.; Conceptualization: S.G.

### Data and materials availability

All raw data are available as supplementary material.

## Materials and Methods

### Protein purification

#### Purification of *E. coli* GF4-3 JetABCD (ecJET)

ecJetABC (WT and variants containing hinge charge-reversal mutations) were purified similarly as previously described ^25^. For ecJetABC lacking native cysteine residues (Cys-less) and its derivatives containing engineered cysteine pairs, the initial purification steps (cell lysis, StrepTrap XT affinity chromatography, 3C tag removal) were performed with buffers supplemented with 5 mM beta-mercaptoethanol. The final size-exclusion chromatography (SEC) was performed with a Superose6 Increase 10/300 GL column equilibrated with Wadjet reaction buffer (ATG buffer, 10 mM HEPES–KOH pH 7.5, 150 mM KOAc, 5 mM MgCl_2_) devoid of any reducing agents. For JetABC variants containing the ecJetC(core) (lacking the elbow-to-hinge module), the same procedure was performed but omitting any reducing agents from the purification process (to allow disulfide bond formation for stable JetC dimerization). The ec-nvJetABC chimera was purified as described in ^25^ for the ecJetABC complex. ecJetD was purified as in ^25^, however, with the last SEC step performed in a column equilibrated in ATG buffer without reducing agents.

#### Purification of *N. vireti* LMG 21834 JetABCD (nvJET)

nvJET was purified as described in ^29^.

#### Reconstitution of Wadjet complexes

Once purified, reaction-ready stocks for all Wadjet variants were reconstituted by mixing ecJetABC (250 nM final dimer, (JetC_2_JetB_2_JetA_1_)_2_ in the case of ecJetABC; (JetC_2_JetB_1_JetA_1_)_2_, in the case of nvJetABC) with JetD (500 nM final dimer, JetD_2_) in either MM buffer (25 mM HEPES-KOH pH 7.5, 250 mM potassium glutamate, 10 mM magnesium acetate), or ATG buffer (10 mM HEPES-KOH pH 7.5, 150 mM KOAc, 5 mM MgCl_2_). Since JetABC subunits are the limiting component of the DNA cleavage reaction, all methods described below will use “ec/nvJET dimer” to refer to a preparation of ec/nvJetABC + ec/nvJetD as detailed above.

#### Purification of *S. cerevisiae* Smc5/6 hexamer

Plasmids containing wild-type or mutant versions of the six core scSmc5/6 subunits were cloned using a recently published procedure ^46^. For all constructs a carboxy-terminal 3C-TwinStrep tag was added, and the complexes were purified following a published protocol ^47^.

#### Purification of *S. cerevisiae* Smc5/6 hinge/coiled coil constructs

Plasmids containing wild type or mutant versions of ybbR-His-tagged scSmc6(407-808) and untagged scSmc5(364-743) were cloned using a recently published procedure ^46^, transformed into chemically competent Rosetta (DE3) cells, and 1 L cultures were grown in TB-medium at 37°C to an OD_600_ of 1. The temperature was then reduced to 22°C and protein production was induced by adding IPTG to a final concentration of 0.4 mM. Cells were incubated overnight (12-16 h), harvested by centrifugation, resuspended in lysis Buffer (50 mM Tris pH 7.5, 300 mM NaCl, 5 % glycerol, 25 mM imidazole, 5 mM beta-mercaptoethanol) and lysed by sonication. The lysate was clarified by centrifugation (40000 *g* for 30 minutes at 4°C) and the supernatant was loaded on a 5 ml HisTrap column (Cytiva). After washing with 10 CV of lysis buffer, the bound material was eluted with a 10 CV gradient from lysis buffer to elution buffer (lysis buffer supplemented with 500 mM imidazole). Peak fractions were analysed by SDS-PAGE, and the fractions with superior purity were concentrated to a final volume of 1 ml, typically yielding concentrations between 15 and 25 mg/ml. 500 μl aliquots were prepared and loaded on a Superdex 200 Increase 10/300 column (Cytiva) in SEC buffer (20 mM Tris pH 7.5, 250 mM NaCl, 1 mM TCEP). Peak fractions were analyzed by SDS-PAGE, pooled, and frozen in aliquots without further purification.

#### Purification of *S. cerevisiae* condensin (Smc2/4) hinge/coiled coil constructs

Budding yeast condensin hinges (scSmc2(443-740), scSmc4(598-923)) were expressed in insect Sf9 cells following the protocol described in ^9^. They were purified by a three strep purification protocol. Cells were lysed by dounce homogenisation and lysates cleared by ultracentrifugation and the lysate collected. Protein was pulled down via its Twin-StrepII tag by incubation with Strep-Tactin XT resin for 2 hours at 4 °C. Protein was eluted by incubation with 100 mM biotin for 30 mins at 4 °C. The tag was cleaved by incubation with 1 mg TEV protease for 2 hours at 4 °C. Protein was then further purified by injection into a cation exchange column and eluted over a gradient of 50 mM to 1 M NaCl. Protein was then injected into a Superdex 200 increase 10/300 GL size exclusion column and the protein peak was pooled, concentrated, and then frozen in liquid N_2_ and stored at – 70 °C.

#### Purification of *B. subtilis* hinge/coiled coil constructs

A construct for the *B. subtilis* hinge/coiled coil (bsSmc(400-776; C437S, R558C, N634C)); ^31^ was modified using standard mutagenesis methods to introduce hinge-channel mutations. All bsSmc hinge/coiled coil expression plasmids were transformed into BL21(DE3) Gold cells, and 1 L cultures of were grown in TB at 37°C to an OD_600_ of 1. The temperature was then reduced to 18°C and protein expression was induced with 0.5 mM IPTG. After overnight (12-16 h) incubation the cells were harvested by centrifugation, and proteins were purified as described for the scSmc5/6 hinge/coiled coil constructs.

#### Purification of *E. coli* GF4-3 JetC hinge/coiled coil constructs

The ecJetC hinge/coiled coil, lacking endogenous cysteine residues but containing engineered cysteine residues for crosslinking, were produced in *E. coli* BL21, expressed with an amino-terminal ybbR-6His tag. A 500 mL culture of the BL21 strain was grown in TB medium at 37°C in a non-baffled glass flask, until the culture reached OD_600_=0.5. The culture was cooled to 16°C and protein overexpression induced by IPTG (0.5 mM final) for 16 hours. Cells were harvested by centrifugation, resuspended in lysis buffer (50 mM Tris pH 7.5, 300 mM NaCl, 5% (v/v) glycerol, 25 mM imidazole, 5 mM beta-mercaptoethanol), followed by lysis via sonication on ice with a VS70T tip using a Bandelin SonoPuls unit, at 40% output for 12 min with pulsing (1 s on / 1 s off). After clarification by ultracentrifugation (40,000 *g* for 30 min), the lysate was loaded onto a HisTrap HP 5 mL column (Cytiva), washed with 6 CV of lysis buffer and eluted with lysis buffer containing an increasing imidazole gradient (up to 500 mM) for 10 CV. Fractions containing ybbR-6His-ecJetC-hinge were collected, concentrated using Amicon Ultracentrifugal filter units (30 kDa cutoff) and injected onto a Superdex 200 Increase 10/300 size-exclusion chromatography (SEC) column, equilibrated with 20 mM Tris–HCl pH 7.5, 250 mM NaCl and 1 mM TCEP. Hinge-containing fractions were pooled, concentrated to about 1 mg/mL (∼12 μM dimer) and flash frozen in liquid nitrogen for long term storage at - 70 C.

### DNA substrate preparation

#### Chemically modified DNA circles

See Table S1 for a list of modified substrates and their starting plasmid/oligo(s). Preparation of biotinylated 2.9 kb DNA circles was done using a previously described plasmid gap-filling method ^48^. 4-5 μg of pSG6970 (pG46, containing one modification site A, ^49^ or pSG7084 (pG46-46, containing two sites A and B) was nicked in rCutsmart buffer with Nt.BbvCI (NEB, 30 U) in 20 μL reactions for one hour at 37°C. The reaction was quenched by addition of 3 μL Tris pH 8 (200 mM) and 3 μL EDTA pH 8 (400 mM). An excess of replacement oligo (1 μL of 100 μM stock) was added to the mix. Nicked ssDNA fragments were melted away from template by heating at 80°C for two minutes, followed by replacement oligo annealing by slowly cooling the reaction to 20°C (1°C decrease every minute). After column purification and elution in 52 μL water, any DNA nicks were sealed by addition of 6 μL T4 ligase buffer and 2 μL T4 ligase (10 Weiss U), followed by incubation for one hour at 22°C. Finally, a second round of column purification was performed.

For doubly modified substrates, the replacement reaction was performed by adding two replacement oligos (1 μL each) into the tube containing nicked pSG7084. To create DNA substrates with dsDNA branches, an excess (3 μL of 100 μM stock) of annealing oligo, fully or partially complementary to the 40 nt ssDNA flap (see Table S1) of STQ15-5-40ssDNAflap replacement oligo added alongside during the annealing step.

#### Preparation of ssDNA circles

Circular ssDNA substrates for entrapment assays for bsSmc, ecJET and scSmc5/6 were prepared as previously described ^50,51^ with minor modifications. A phagemid based on pBluescript SK-(pDHJS4 AS-; ^51^) was obtained from Addgene (#78243) and transformed into competent *E. coli* DH5a cells carrying a helper plasmid (HP4_M13 ^52^) (Addgene #120340). Cells were plated on LB plates containing 100 μg/ml ampicillin and 50 μg/ml kanamycin, and five individual colonies were inoculated overnight (12-16 h) in 10 ml TB medium containing both antibiotics. Cultures with cells in exponential phase were then used to inoculate 3 L of the same medium in glass flasks without baffles. 5 L flasks were used but only filled with 500 ml of medium to ensure sufficient aeration. The flasks were incubated until the next morning at 37°C shaking at 120 rpm. Bacterial cells were pelleted at 8000 *g* for 20 minutes at 4°C, and the supernatant was collected and subjected to a second round of centrifugation to remove as many bacterial cells as possible. The supernatant was collected in a beaker, cooled to 4°C, and ¼ volume of 5 x phage precipitation buffer (25 % PEG 6000, 2.5 M NaCl) was slowly added while stirring the liquid with a magnetic stir bar. Precipitation continued overnight at 4°C, and phages were then collected by centrifugation at 12000 *g*. The obtained pellet was then resuspended in phage lysis Buffer (10 mM MOPS pH 7.5, 500 mM guanidine-HCl, 1 % Triton X-100) and heated to 80°C in a water bath for 45 min, with gentle mixing by inversion every 5 minutes. After cooling to room temperature, the obtained solution was loaded onto a column from the NucleoBond Extra Maxi Kit (Macherey-Nagel) and DNA was washed, eluted, and precipitated according to manufacturer instructions.

For the entrapment assay with scSmc2/4, the same ssDNA circles were used. pBluescript SK-first transformed into JM109 *E. coli* cells. A colony was picked and inoculated in 1 L 2x TY media plus appropriate antibiotic and cells were grown at 37 °C 200 rpm until OD_600_ = 0.05. M13K07 phage was then added, and cells were grown for a further 90 min. Kanamycin was then added to a concentration of 70 µg/ml and cultures grown overnight. Cells were pelleted and the supernatant was collected. Phages were precipitated by adding 40 g PEG 6000 and 29.2 g NaCl and incubated at 4 °C for 90 mins. Phages were pelleted by centrifugation, washed with 1x TE buffer and then pelleted again. The phage pellet was collected and resuspended in CsCl solution of a density of 1.35 g/ml. The solution was then centrifuged in an SW41 rotor at 280,000 g at 4 °C for 24 hours. The phage content was extracted and ssDNA purified by phenol chloroform extraction followed by dialysis against 50 mM HEPES pH 7.5.

### Cleavage assay with ecJET on obstacle-free DNA

The cleavage assay performed with M4M-crosslinked Wadjets on obstacle-free bio40flapDNA (Extended Data Fig. 3D) was conducted in 15 μL (per reaction condition) MM buffer (25 mM HEPES pH 7.5, 250 mM potassium glutamate, 10 mM magnesium acetate) supplemented with ATP (1 mM final). Typically, ∼7 nM DNA was mixed with ∼15 nM ecJET dimer and/or ScaI (NEB, 10 U) for 15 minutes at 37°C (per incubation step). To remove salt that hindered DNA migration and visualization in agarose gels, the completed reactions were column purified into 15 μL water. This was followed by addition of 3 μL purple loading dye (NEB). The reactions were loaded onto an EtBr-containing 1% (w/v) agarose gel, ran at 5 V/cm for 1 hour and bands visualized with a transilluminator (UVP GelSolo).

The cleavage assay performed with ecJET(core) on pDonor (Extended Data Fig. 4A) was performed in 15 μL ATG buffer (10 mM HEPES-KOH pH 7.5, 150 mM KOAc, 5 mM MgCl_2_) supplemented with 1 mM ATP, containing 8.5 nM pDonor and 12.5 nM ecJET dimer. 2 mM final DTT was added as indicated. Reactions were incubated for 15 minutes at 37°C, followed by addition of 3 μL 6 x SDS-containing loading dye (Thermo Fisher Scientific) and heating at 70°C for 10 minutes. The reactions were loaded onto an EtBr-containing 1% (w/v) agarose gel, ran at 5 V/cm for 1 hour and bands visualized with a transilluminator (UVP GelSolo).

### Experiments with DNA substrates anchored to streptavidin-coated Dynabeads

#### Cleavage assay with ecJET and nvJET

Cleavage assays on Dynabead-bound DNA were performed as described previously ^25^ with minor modifications. Per reaction condition, 10 μL (containing 100 μg) Dynabeads™ MyOne™ Streptavidin C1 were equilibrated and washed with 1xBW buffer (5 mM Tris-HCl pH 7.5, 0.5 mM EDTA, 1 M NaCl) according to manufacturer’s protocol. The beads were resuspended in 20 μL 2xBW buffer (10 mM Tris-HCl pH 7.5, 1 mM EDTA, 2 M NaCl) and incubated with an equal volume of biotinylated DNA (200 ng in water per condition) at 30°C for 30 minutes with shaking. After washing with 1xBW buffer, the beads were equilibrated with MM reaction buffer (25 mM HEPES pH 7.5, 250 mM potassium glutamate, 10 mM magnesium acetate).

Equilibrated beads were then resuspended in MM buffer supplemented with 1 mM ATP (15 μL per reaction, scaled accordingly). For experiments involving nvJET, 5 mM final MnCl_2_ was supplemented to the reaction buffer. For experiments involving multiple conditions, this was split into 15 μL aliquots. Beads were treated with Wadjet (12.5 nM dimerfor ecJET either freshly reconstituted or crosslinked with M4M as described below, 25 nM dimer for nvJET) and/or ScaI-HF (NEB, 10 U), 15 minutes at 37°C per incubation step. DNA bound to the beads was eluted by addition of 85 μL preheated MM buffer supplemented with 25 mM biotin and SDS (0.1% final) and incubation for 10 minutes at 70°C. The supernatant containing the eluate was separated from the beads with a magnetic rack. To remove salt that hindered DNA migration and visualization in agarose gels, the eluted DNA was column purified into 15 μL water. This was followed by addition of 3 μL purple loading dye (NEB). The reactions were loaded onto an EtBr-containing 1% (w/v) agarose gel, ran at 5 V/cm for 1 hour and bands visualized with a transilluminator (UVP GelSolo).

#### DNA leakage assay

To determine the stability of biotinylated DNA circles on streptavidin-coated Dynabeads (DNA leakage), DNA was bound to Dynabeads™ MyOne™ Streptavidin C1 as above, and after washing and equilibration, the beads were resuspended in 15 μL MM buffer and incubated at 37°C for 30 minutes. The supernatant containing the released fraction was isolated. The beads containing still-bound material was eluted by the addition of 85 μL preheated MM buffer supplemented with 25 mM biotin and SDS (0.1% final) for 10 minutes at 70°C. The supernatant containing the elution fraction was collected from the beads. Both fractions were column purified as above and loaded onto an EtBr gel.

### *in vitro* protein crosslinking

#### Preparative cysteine crosslinking with 1,4-butanediyl bismethanethiosulfonate (M4M)

Purified ecJetABC (Cys-less and its derivatives) were diluted to 5 μM dimer in ATG buffer to a volume of 25 μL. Crosslinking proceeded by adding 1.25 μL of 1,4-butanediyl bismethanethiosulfonate (M4M) (diluted to 2 mM in DMSO, LGC Standards, 100 μM final) to the protein, and the reaction was left overnight at 4°C. The crosslinked protein solution was spun down (21 000 *g*) for 10 minutes at 4°C to remove any aggregate, its concentration was determined and subsequently diluted to 250 nM dimer in MM buffer supplemented either with quenchers *S*-Methyl methanethiosulfonate (MMTS, Merck, 5 mM final preserving the crosslinks) or with dithiothreitol (DTT, BioChemica, 5 mM final disrupting the crosslinks). The quenching took place in room temperature for 30 minutes, before transfer onto ice. ecJetD (500 nM dimer final) was added to the quenched protein to produce a reaction-ready preparation.

#### Analytical crosslinking of SMC hinge/coiled coil constructs with BMOE

Crosslinking reactions for bsSmc, scSmc5/6 and ecJET were conducted in 10 μL crosslinking buffer (20 mM Tris, 50 mM NaCl). ecJetC hinges (0.5 μM dimer final) were mixed with/without 0.5 μL of BMOE (20 mM stock, 1 mM final) for 45 seconds before quenching by addition of 1 μL DTT (100 mM stock). The crosslinked reactions were analyzed via SDS-PAGE.

For scSmc2/4, protein was diluted to 1.4 μM final in CB buffer (50 mM NaCl, 1 mM MgCl_2_, 50 mM HEPES pH 7.5, 1 mM TCEP and 5 % glycerol) to a final volume of 10 µl. To this, either 1 µl DMSO or 1 µl of BMOE (6.4 mM stock, 0.58 mM final) was added and mixtures incubated on ice for 6 min. 4 x LDS buffer (ThermoFisher) was then added and reactions heated at 70 °C for 10 min. Reaction products were then separated by SDS-PAGE.

### DNA entrapment assays with isolated hinge/coiled coil constructs

For the entrapment assays with bsSmc, scSmc5/6 and ecJET hinge/coiled coils, 30 μl reactions were set up containing 30 nM ssDNA or 15 nM dsDNA circles and 500 nM of protein complex in a buffer composed of 10 mM Tris-HCl pH 7.5 and 50 mM NaCl. The mixture was incubated at RT for 5 minutes before 25 μl were transferred to a fresh 1.5 mL Eppendorf tube containing 1.25 μl of 20 mM BMOE (final concentration 1 mM). After 45 seconds of incubation at room temperature, 25 μl were transferred again to another fresh 1.5 mL Eppendorf tube containing 2.5 μl of 100 mM DTT for quenching (final DTT concentration 10 mM). 5 μl of 6 x loading dye with SDS (ThermoFisher) were added and the samples were heated to 70°C for 10 minutes to denature the proteins. Aliquots of 10 μl were then separated on 1 % agarose gels (in 0.5x TBE) containing 0.03 % SDS (Sigma-Aldrich) and ran in 0.5x TBE buffer with 0.03 % SDS at 7.5 V/cm for about 1 hour at room temperature.

For the entrapment assay with scSmc2/4 hinge/coiled coils, protein was diluted to a concentration of 325 nM and ssDNA to 9.3 nM in EB buffer (20 mM NaCl, 1 mM MgCl_2_, 50 mM HEPES pH 7.5, 1 mM TCEP and 5 % glycerol) in a volume of 13 μl. Samples were incubated at 24 °C for 60 min. To these, either 1 μl DMSO or 1 μl of 6.4 mM BMOE was added and mixtures incubated on ice for 6 min. 6 x DNA loading dye (NEB) was added and mixtures heated at 70 °C for 20 min. Reaction products were then separated by electrophoresis with 0.8 % agarose gels ran at 100 V for 4 hours in 1x TBE.

### DNA entrapment assays with hexameric Smc5/6 complexes

Crosslinking reactions were set up as described for hinge/coiled coil constructs, with the exception that a modified buffer (20 mM Tris-HCl pH 7.5, 150 mM NaCl, 20 % glycerol) was used due to reduced protein stability of the hexameric core complex under low salt conditions.

### Strain construction in *E. coli*

Integration plasmids (listed in Table S2) containing wild-type or hinge mutant *E. coli* GF4-3 *jetABCD* under the arabinose-inducible pBAD promoter were cloned into MFDpir *E. coli* donor. Tri-parental mating between the donor and *E. coli* K12 recipients was performed as previously described ^53^, resulting in insertion of the *jet* operons via transposon mediated integration into the recipient genome near the neutral chromosomal *glmS* loci. The test plasmid pBADMycHisA (pBAD, 4 kb) was introduced to these strains via electroporation at 2.0 kV. See Table S3 for a list of bacterial strains used.

### Plasmid stability assay in *E. coli*

Plasmid stability assays were performed as described ^25^. Briefly, pBAD and *jet* operon containing *E. coli* strains were grown overnight in LB media in the presence of ampicillin (100 μg/mL final), they were backdiluted 1000-fold in fresh LB media without antibiotics but containing arabinose (0.02% w/v final). After 6-7 hours when cells have undergone approximately ten generations of growth, they were spotted in 10-fold dilution series on either nutrient agar (NA) plates (scoring for overall cell number), or NA plates containing ampicillin (100 μg/mL final, scoring for ampicillin-resistant plasmid-containing survivors). After overnight incubation at 37°C, colonies were counted and extrapolated based on the dilution degree to give an estimate of the total cell number and survivor count. To quantify pBAD+ cells, the percentage of ampicillin-resistant cells over the total cell number was calculated.

### Strain construction in *B. subtilis*

Transformation with recombinant DNA was used to engineer *B. subtilis* strains at the *smc* loci by allelic replacement using starvation-induced natural competence under standard conditions (1-2 h incubation in SMM medium lacking amino acid supplements) as described in ^54^. Strains were selected on SMG-agar plates under appropriate antibiotic selection at 37°C. Genotypes were verified for selected single colony isolates by antibiotic resistance profiling, colony PCR, and Sanger sequencing as required. See Table S3 for a list of bacterial strains used.

### Viability assessment by dilution spotting

*B. subtilis c*ultures were inoculated in SMG medium and grown for 10 h at 37°C under constant shaking. Cultures were diluted 1:9 in series. Dilutions of 9^2^ and 9^5^ were spotted on minimal SMG agar and nutrient agar (ONA) plates and grown at 37°C. Colony growth was documented by imaging after 16 h for NA plates and 24 h for SMG-agar plates.

### Chromatin immunoprecipitation (ChIP)

ChIP samples were prepared as described previously ^55^. Cultures were grown in 200 mL SMG at 37°C. Cells were grown to mid-exponential phase (OD_600_ = 0.02–0.03) and fixed by incubation for 30 min with 20 mL buffer F (50 mM Tris-HCl pH 7.4, 100 mM NaCl, 0.5 mM EGTA pH 8.0, 1 mM EDTA pH 8.0, 10% (w/v) formaldehyde). Cells were harvested by filtration and washed in cold PBS. The OD_600_ values of the samples were normalized to 2 and resuspended in TSEMS (50 mM Tris pH 7.4, 50 mM NaCl, 10 mM EDTA pH 8.0, 0.5 M sucrose and protease inhibitor cocktail (PIC) (Sigma)) supplemented with 6 mg/mL chicken egg white lysozyme (Sigma). Samples were incubated at 37°C for 30 min under rigorous shaking. The resulting protoplasts were harvested by centrifugation, washed in 1 mL TSEMS, resuspended in 1 ml TSEMS and split into 3 aliquots of equivalent volume before pelleting and flash freezing.

Samples were resuspended in 2 mL of buffer L (50 mM HEPES-KOH pH 7.5, 140 mM NaCl, 1 mM EDTA pH 8.0, 1% (v/v) Triton X-100, 0.1% (w/v) Na-deoxycholate, 0.1 mg/mL RNaseA and PIC (Sigma)), transferred to 5 mL round-bottom tubes and sonicated three 20 second pulses using a Bandelin Sonoplus with an MS72 tip (90% pulse and 35% power output). Suspensions were centrifuged for 10 minutes at 21000 *g* at 4°C. Samples were split into 200 µL input material and 800 µL IP material. Anti-ScpB antibody serum (Eurogentec) was incubated with equivalent volumes of Dynabeads Protein G suspension (Invitrogen) for 2 h at 4°C under gentle agitation. Beads were washed in 1 mL Buffer L directly prior to use and resuspended as 50 µL aliquots. IP material was mixed with these 50 µL aliquots and incubated at 4°C for 2 hours under rotation.

Bound material was subsequently washed by 1 mL washes with buffer L, L5 (buffer L containing 500 mM NaCl), buffer W (10 mM Tris-HCl pH 8.0, 250 mM LiCl, 0.5% (v/v) NP-40, 0.5% (w/v) Na-deoxycholate, 1 mM EDTA pH 8.0) and buffer TE (10 mM Tris-HCl pH 8.0, 1 mM EDTA pH 8.0). Beads were resuspended in 520 µL TES (50 mM Tris-HCl pH 8.0, 10 mM EDTA pH 8.0, 1% (w/v) SDS). Input material was supplemented with 300 µL TES and 20 µL 10% SDS. Tubes were incubated at 65°C overnight under vigorous shaking. DNA was purified using two rounds of phenol-chloroform extraction. 400 µL of extracted DNA was mixed with 1.2 µL GlycoBlue (Invitrogen), 30 µL of 3 M Na-acetate (pH 5.2) and 1 mL ethanol (filtered). Samples were incubated at -20°C for 20 minutes. Precipitated DNA was pelleted by centrifugation at room temperature at 21000 *g* for 10 minutes and was dissolved by incubation at 55°C in 100 µL EB (Qiagen) under vigorous shaking for 10 minutes. Samples were subsequently purified using a PCR purification kit (Qiagen) as per protocol and eluted in 50 µL EB.

For quantification by qPCR, samples were diluted to 1/10 for IP and 1/1000 for input material. Reactions for qPCR were prepared by mixing 4 µL diluted samples with 5 µL 2 x 5 µL Takyon SYBR MasterMix and 1 µL qPCR primer mixture (3 µM). A list of qPCR primers is given in Table S1. Samples were run in a Rotor-Gene Q (Qiagen) and analyzed using PCR miner ^56^.

