## Extended Data Figures and Legends for "The SMC Hinge is a Selective Gate for Obstacle Bypass"

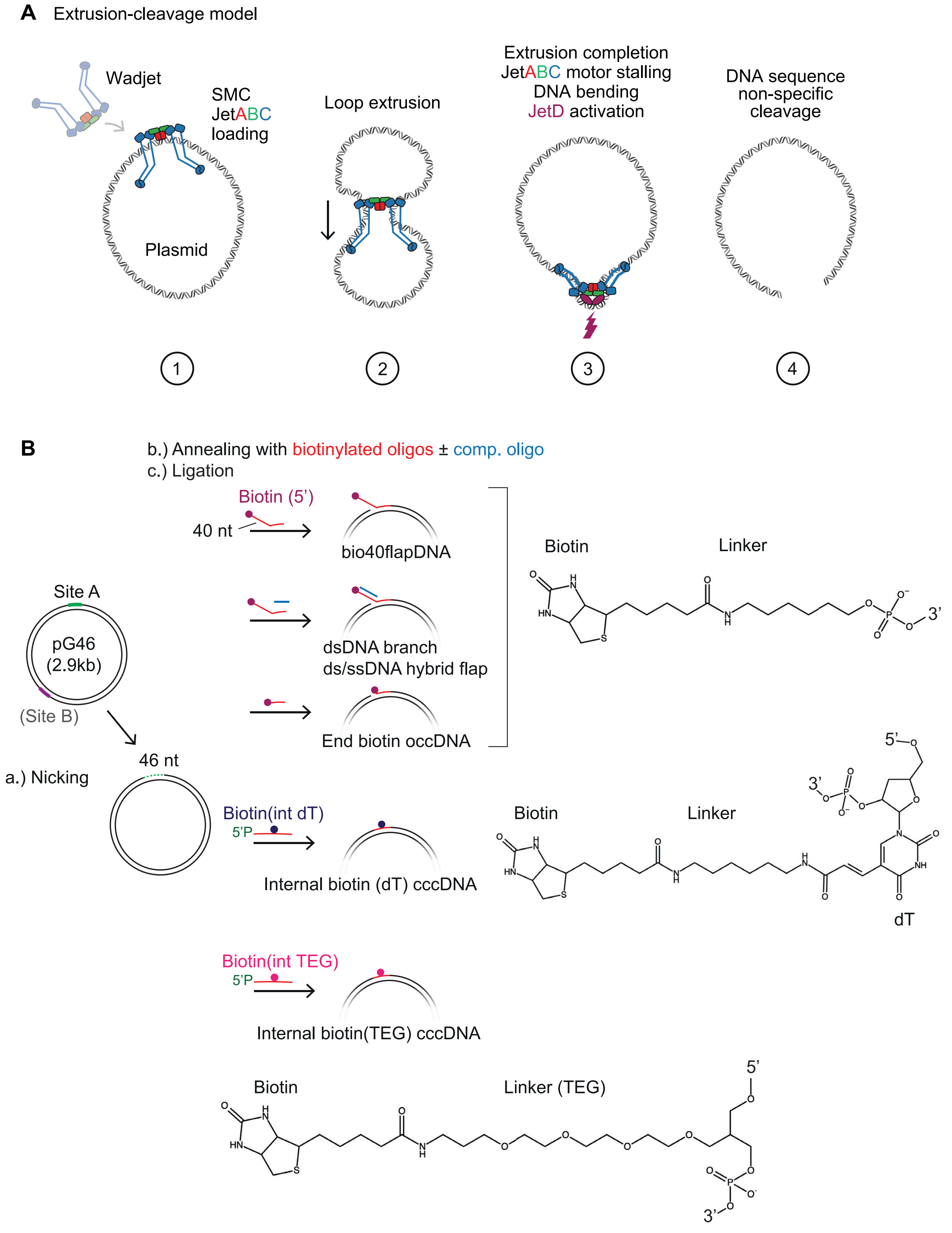


### Extended Data Fig. 1

**Supplementary figure legends**

Extended Data Fig. 1 – Schematics of plasmid restriction by Wadjet and DNA substrate generation.

A) Schematic depicting the extrusion-cleavage model for plasmid restriction by SMC Wadjet. The sensor SMC motor component JetABC loads onto plasmid (step 1), followed by bidirectional DNA extrusion (step 2). On smaller circular substrates, extrusion completion results in JetABC motor stalling, forming the cleavage competent state and mechanical DNA bending [25], triggering JetD activation (step 3) and subsequent DNA sequence non-specific cleavage (step 4).

B) Schematic depicting generation of biotinylated DNA circles used in this study. pG46 [49] containing closely spaced BbvCI sites (site A) was treated with Nt.BbvCI nicking enzyme, which results in short ssDNA fragments that can be molten away upon heating to produce a 46 nt ssDNA gap. Annealing with an excess of replacement oligonucleotide (in red) and subsequent ligation (nick sealing) will generate the indicated biotinylated species. To generate circles with a dsDNA-ssDNA hybrid flap or a dsDNA branch, an excess (relative to the replacement oligonucleotide) of (partially or fully respectively) complementary oligonucleotide (in blue) was added during the annealing step. For circles with two modifications, a pG46 derivative pG46-46 [25]containing an extra Nt.BbvCI-nicking cassette (site B) was used. occDNA - open covalent circles (containing nicks); cccDNA - closed covalent circles (containing no nicks). Chemical structures of the various organic linkers between the biotin moiety and DNA are also displayed. Note that DNA containing internal biotin (dT) contains a biotin moiety attached via a carbon chain to a defined thymidine (dT) nucleobase, while circles containing internal biotin (TEG) contains biotin attached via a longer TEG (tetraethylene glycol) linker between two nucleobases.  See Methods for more details.

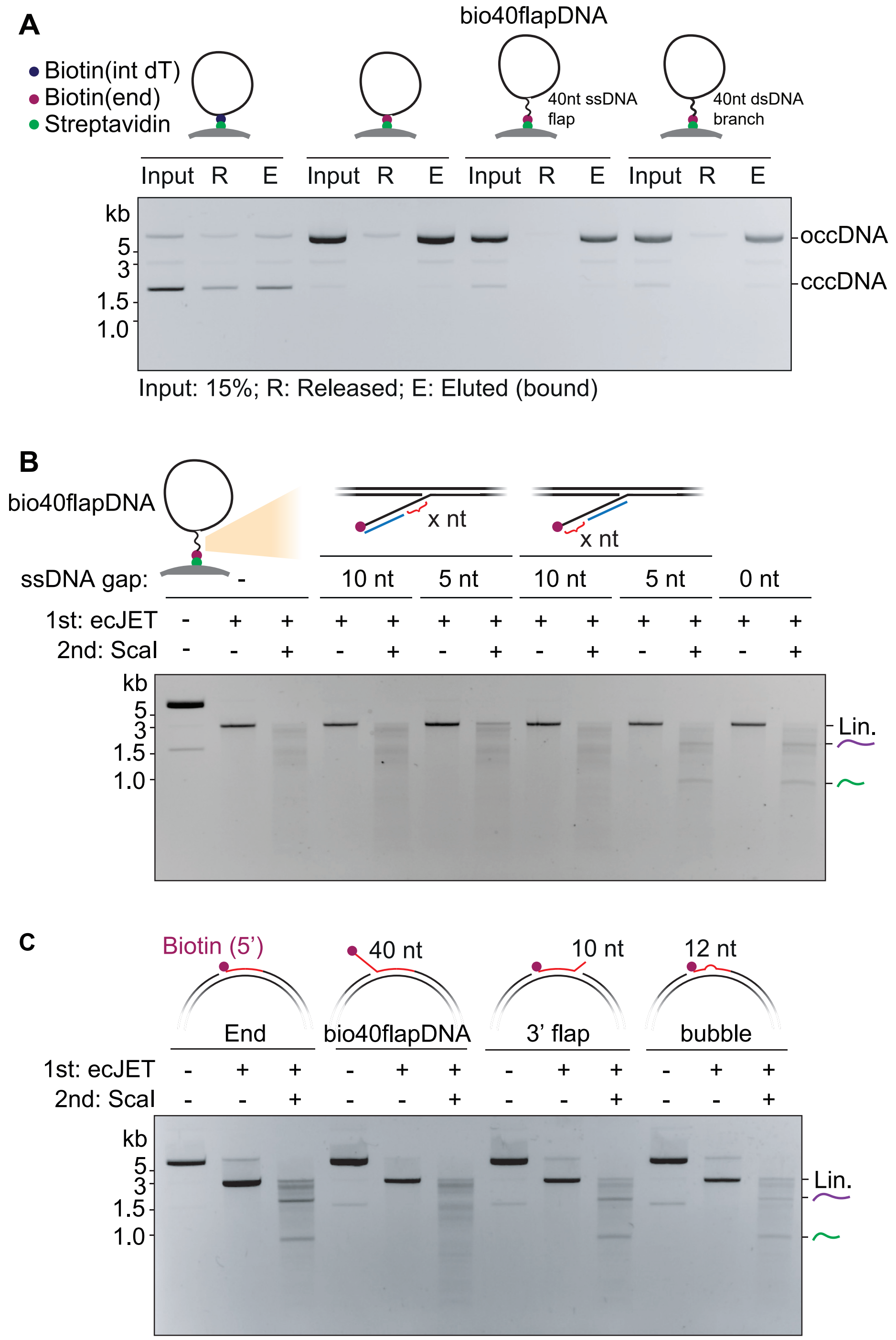


### Extended Data Fig. 2

Extended Data Fig. 2 – Characterization of ssDNA-linked obstacle bypass by Wadjet-I.

A) Agarose gel depicting the propensity for dissociation (leakage) of biotinylated DNA from streptavidin-coated Dynabeads. Beads containing coupled DNA were mock treated in Wadjet reaction buffer for 30 minutes. The supernatant fraction containing released (R) DNA was harvested. The remaining bound DNA was eluted as the elution fraction (E, see Methods). Note that internally biotinylated circles had a higher tendency to leak than 5'-end-biotinylated circles, as observed previously [25] . Presence of a ssDNA flap or a dsDNA branch did not influence the degree of leakage. cccDNA: Closed covalent circular DNA; occDNA: open covalent circular DNA (nicked).

B) Agarose gel showing the cleavage activity of ecJET on bio40flapDNA attached onto Dynabeads, with a partially complementary oligo (in blue) annealing at the ssDNA flap (either towards the biotin end or the junction end), leaving behind defined stretches of ssDNA at the un-annealed end.

C) Agarose gel showing the cleavage activity of ecJET on the indicated biotinylated DNA circles containing ssDNA not constituting the obstacle linker.

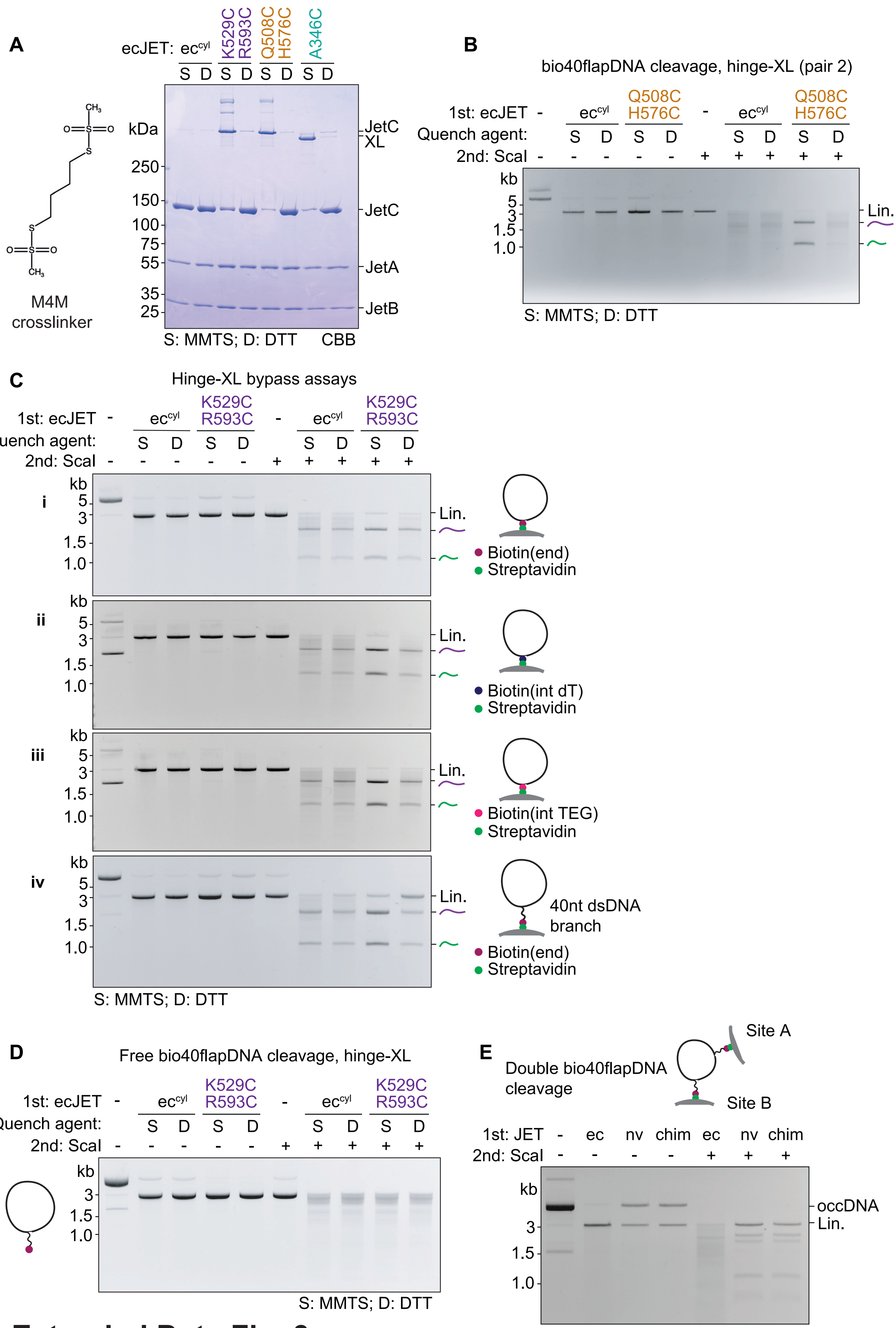


**Extended Data Fig. 3**

Extended Data Fig. 3 - Characterization of hinge bypass.

A) Left: Chemical structure of the 1,4-butanediyl bismethanethiosulfonate (M4M) crosslinker used. Right: SDS-PAGE of Cys-less ecJET derivatives containing engineered cysteines at hinge (pair 1: R529C, R593C; pair 2: Q508C, H576C) or at elbow positions (A346C) after M4M treatment and subsequent quencher treatment (see Methods) - S: MMTS (non-reducing) resulting in slower migrating (crosslinked) species; D: DTT (reducing), resulting in almost complete reversal of crosslinking.
B) Agarose gel depicting the cleavage/bypass activity on bio40flapDNA attached onto Dynabeads by Cys-less ecJET variants, after M4M-induced cysteine crosslinking (of the hinge domain at Q508C, H576C as indicated) and subsequent quenching by the MMTS quencher (preserving the crosslinking) or DTT (disrupting the crosslinking).
C) Agarose gel depicting the cleavage/bypass activity by hinge-crosslinked ecJET on DNA substrates attached onto Dynabeads in different ways: i) circles containing a 5' end-positioned biotin, ii) circles containing internal biotin (dT), iii) circles containing internal biotin (TEG), iv) circles containing 40 bp dsDNA branches. All substrates were treated with Cys-less-ecJET variants, after M4M-induced cysteine crosslinking (of the hinge domain at K529C, R593C) and subsequent quenching by the MMTS quencher (preserving the crosslinking) or DTT (disrupting the crosslinking).
D) As with C) but on bio40flapDNA without Dynabead presence (in solution).
E) Agarose gel showing the cleavage activity of the indicated Wadjet variants on DNA circles with two bio40flap-Dynabead modifications. Wadjets containing a nvJET-hinge cleaved these substrates less efficiently. Residual cleavage events were likely due to a population of DNA circles that have only attached onto a streptavidin Dynabead at one of the two sites.

A

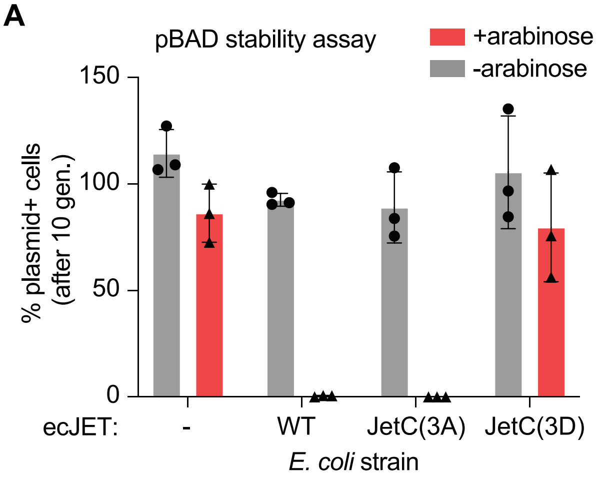


B

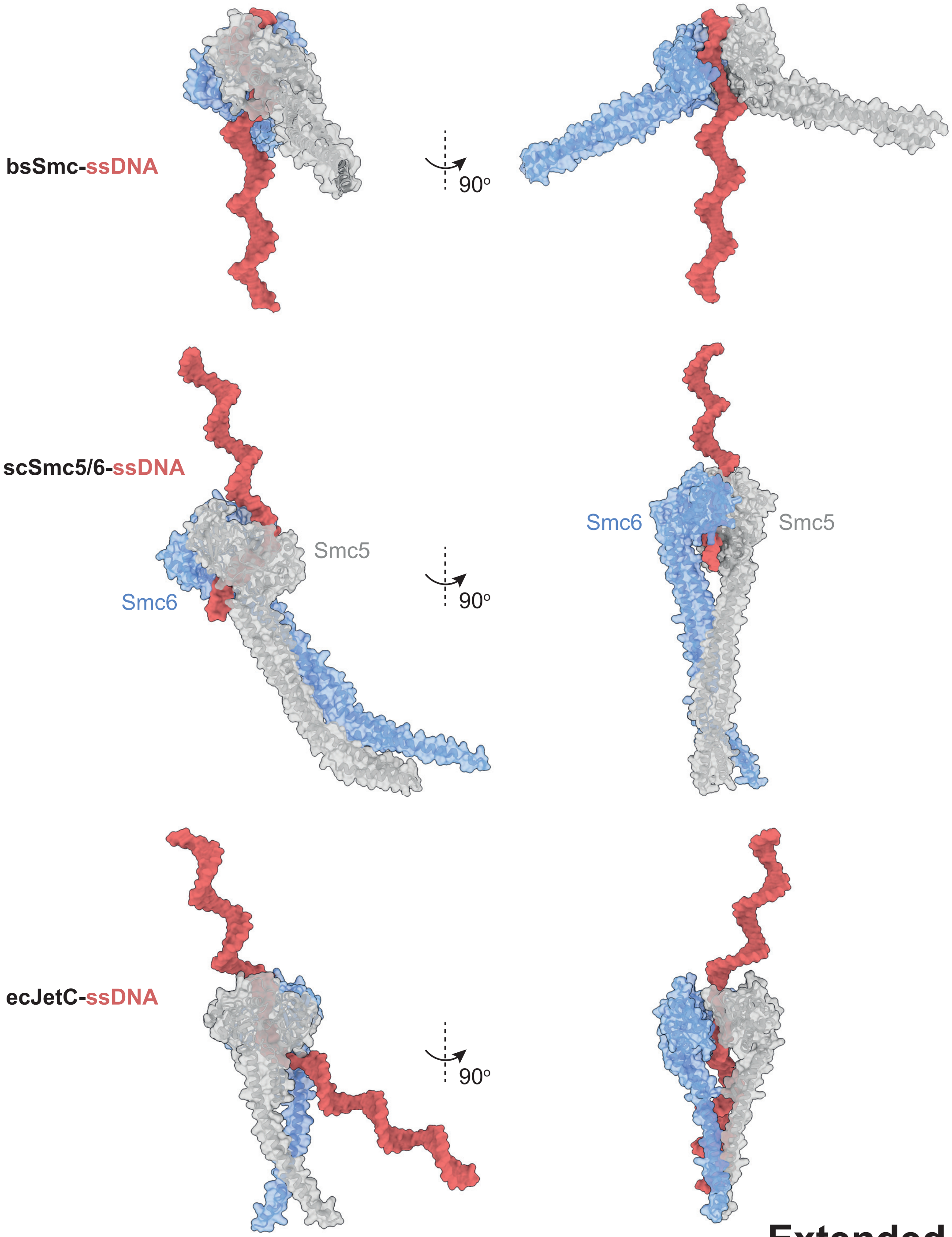


**Extended Data Fig. 4**

Extended Data Fig. 4 - Characterization of ecJET(core) and JetC hinge bypass-deficient mutants.

A) Graph showing the percentage of plasmid (pBAD, 4 kb, AmpR)-containing *E. coli* cells in a cell population grown for ten generations without antibiotic selection with or without induction of the indicated ecJET hinge mutants by arabinose addition. 3A: ecJetC(R524A, K591A, K601A); 3D: ecJetC(R524D, K591D, K601D). Means, standard deviations, and individual data points from three independent experiments are shown. Datapoints greater than 100% were sometimes obtained if the number of plasmid+ colony count was greater than total cell colony count.

B) AlphaFold3 predictions of selected SMC hinge/coiled coils with ssDNA. ecJET, bsSmc, and scSmc5/6 hinge/coiled coil sequences were predicted as dimers together with a stretch of ssDNA (between 36 and 64 nt long), frequently displaying the ssDNA molecule inside a closed or half-open hinge channel. Selected models are shown as representative examples.

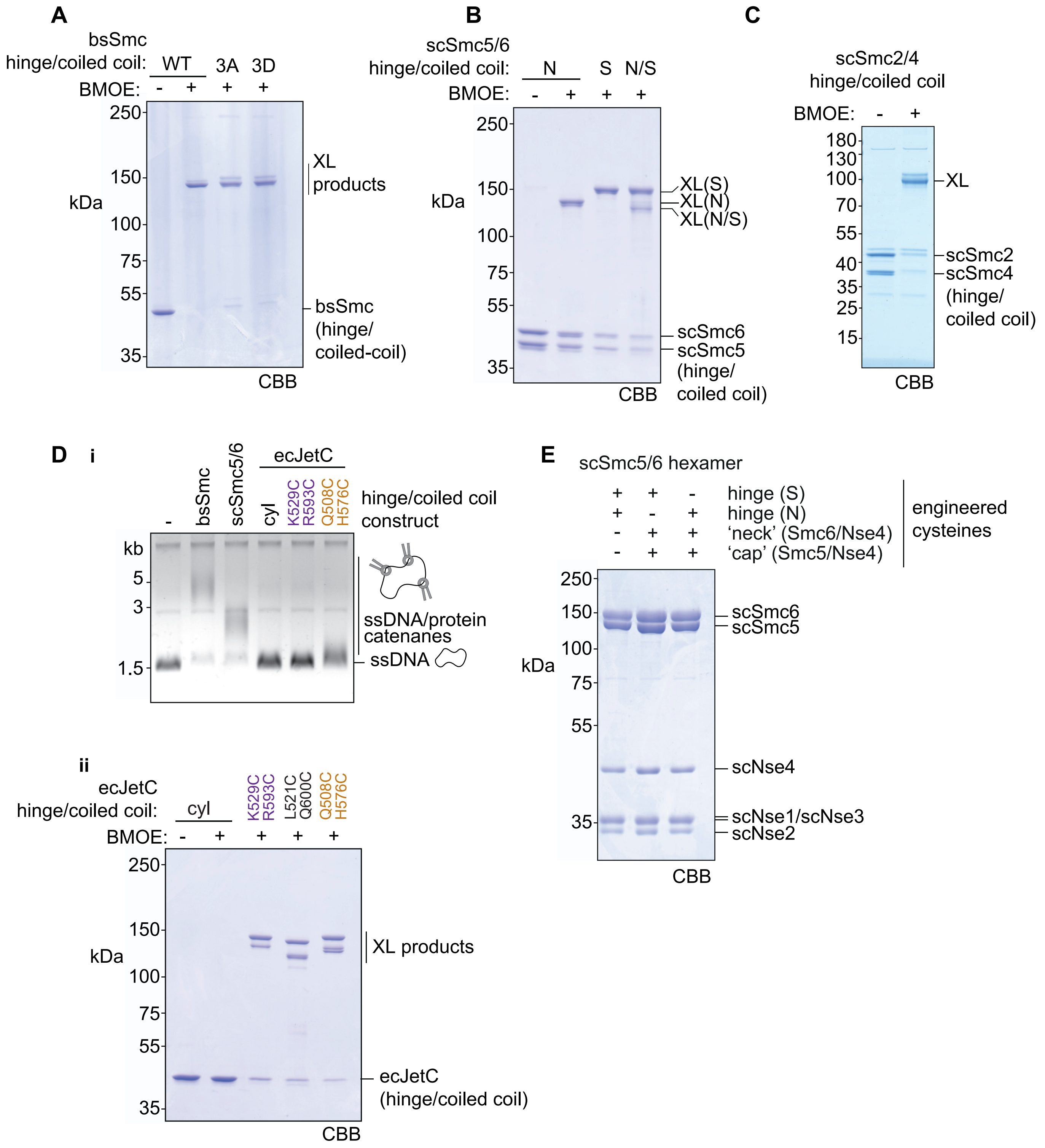


### Extended Data Fig. 5

Extended Data Fig. 5 - Characterisation of a ssDNA-entrapment by SMC complexes.

A) SDS-PAGE showing BMOE crosslinking for the indicated bsSmc hinge/coiled coil constructs (residues 400-776). All hinges harbor C437S to mitigate non-specific crosslinking as well as engineered cysteines R558C and N634C for crosslinking by BMOE. 3A: bsSmc(R558C, N634C, R583A, R643A, R645A); 3D: bsSmc(R558C, N634C, R583D, R643D, R645D).

B) SDS-PAGE showing BMOE crosslinking for the indicated scSmc5/6 hinge/coiled coil constructs (Smc5 residues 305-805, Smc6 residues 407-808). N: “North” interface containing Smc5(V638C), Smc6(N572C); S: “South” interface containing Smc5(N526C), Smc6(N643C).

C) SDS-PAGE showing BMOE crosslinking for the scSmc2/4 hinge/coiled coil construct (Smc2 residues 443-740, Smc4 residues 598-923), containing residues Smc2(S560C, K639C), Smc4(V721C, M821C) for crosslinking.

D) i. Denaturing agarose gel depicting 2.3 kb circular ssDNA entrapment by the indicated ecJetC hinge/coiled coil constructs. bsSmc and scSmc5/6 hinge/coiled coils were included as positive controls. Note that the cysteine pairs used for crosslinking here (with BMOE) were also used for the bypass experiments (crosslinking with M4M). Only the Q508C, H576C pair exhibited modest entrapment. ii. SDS-PAGE showing BMOE crosslinking for the indicated ecJetC hinge/coiled coil constructs. The multiple XL products are a mixture of crosslinks between one (or the other or both) cysteine pair interfaces indistinguishable by mere inspection. Indeed, we suspect the inefficiency of ssDNA entrapment by the ecJetC hinges in our assay might have arisen from inefficient covalent closure of the hinge donut (requiring both cysteine pairs to be crosslinked) due to the relatively short spacer between the reactive maleimide moieties of BMOE.

E) SDS-PAGE showing the intactness of scSmc5/6 hexamer constructs, containing the indicated engineered cysteines for crosslinking that were used for ssDNA entrapment assays in Fig. 3Dii.

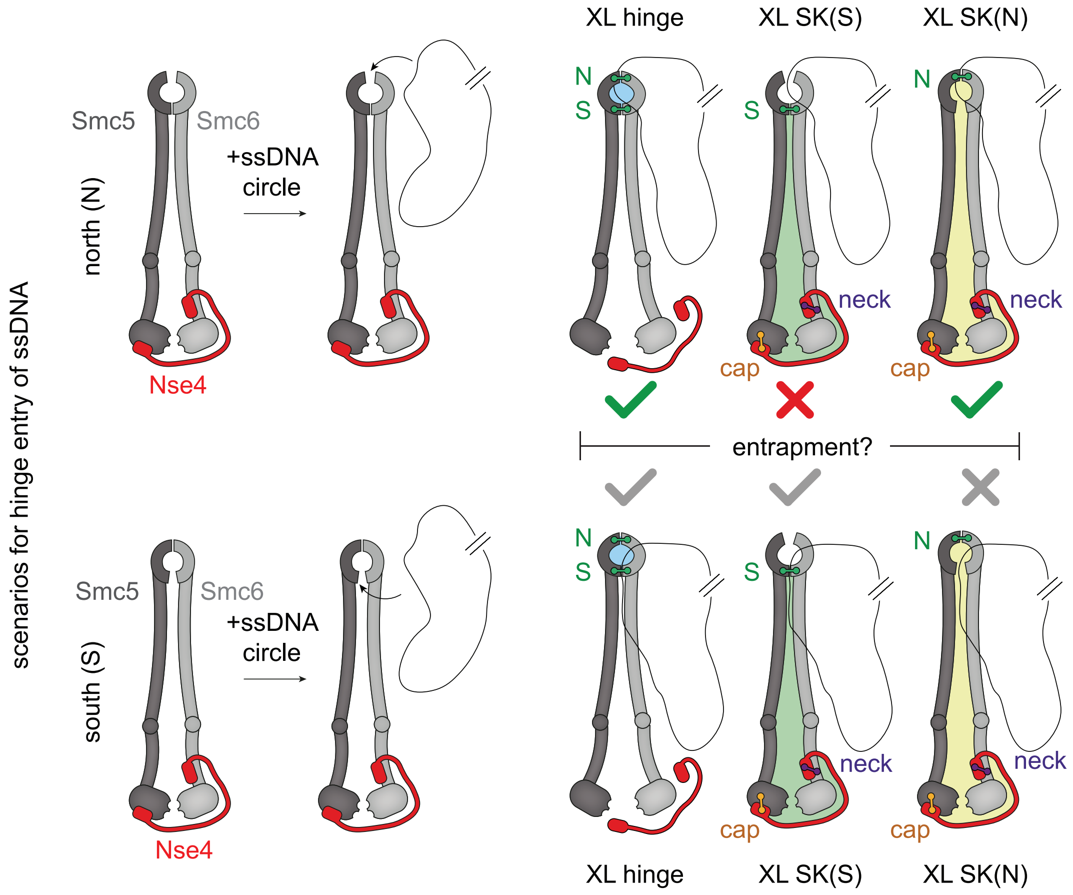


### Extended Data Fig. 6

Extended Data Fig. 6 - ssDNA trajectory into the Smc5/6 ring.

Schematic depiction of the two possible scenarios for ssDNA entry into the hinge channel, via the North (top panels) and the South (bottom panels) interface. Interface crosslinks are depicted as colored dumbbells, covalently closed compartments are indicated by colored areas. Whereas crosslinking of both hinge-interfaces leads to stable entrapment in both cases, combination of individual hinge-interfaces with both Smc/kleisin ('cap' and 'neck') interfaces can distinguish between the scenarios. Upon entry via the North interface, the resulting SMC/kleisin (SK) compartment only contains DNA if the hinge is closed at the North interface. Upon entry via the South interface, the opposite is the case. Note that even though DNA is present in the SK(N) compartment, it completes two passages and thus is pseudotopologically bound. Only the top scenario matches the experimental observation (Fig. 3Dii).

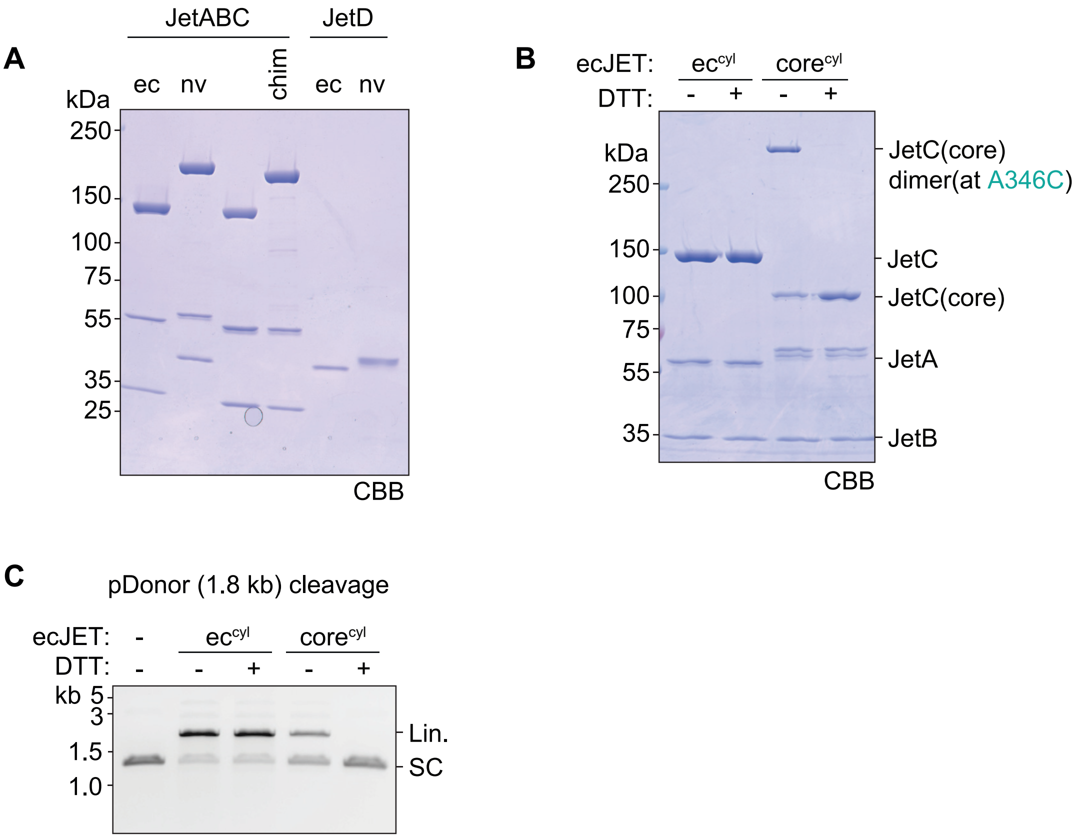


**Extended Data Fig. 7**

Extended Data Fig. 7 – Purification and characterization of engineered ecJET variants

A) SDS-PAGE showing the migration profiles of the indicated Wadjet proteins.
B) SDS-PAGE showing the dimerization of Cys-less ecJetC(core) in non-reducing conditions, through the disulfide bonds between the engineered A346C residues (naturally formed during purification sans reducing agents). DTT presence as expected reduced and de-crosslinked these bonds.
C) Agarose gel showing the cleavage activity of the indicated Wadjet variants on pDonor (1.8 kb). DTT (2 mM final) was added as indicated. We note that cleavage activity of Cys-less ecJET(core) is deficient compared to WT, and that DTT addition notably abrogated cleavage activity of ecJET(core) due to loss of complex integrity. SC: supercoiled.

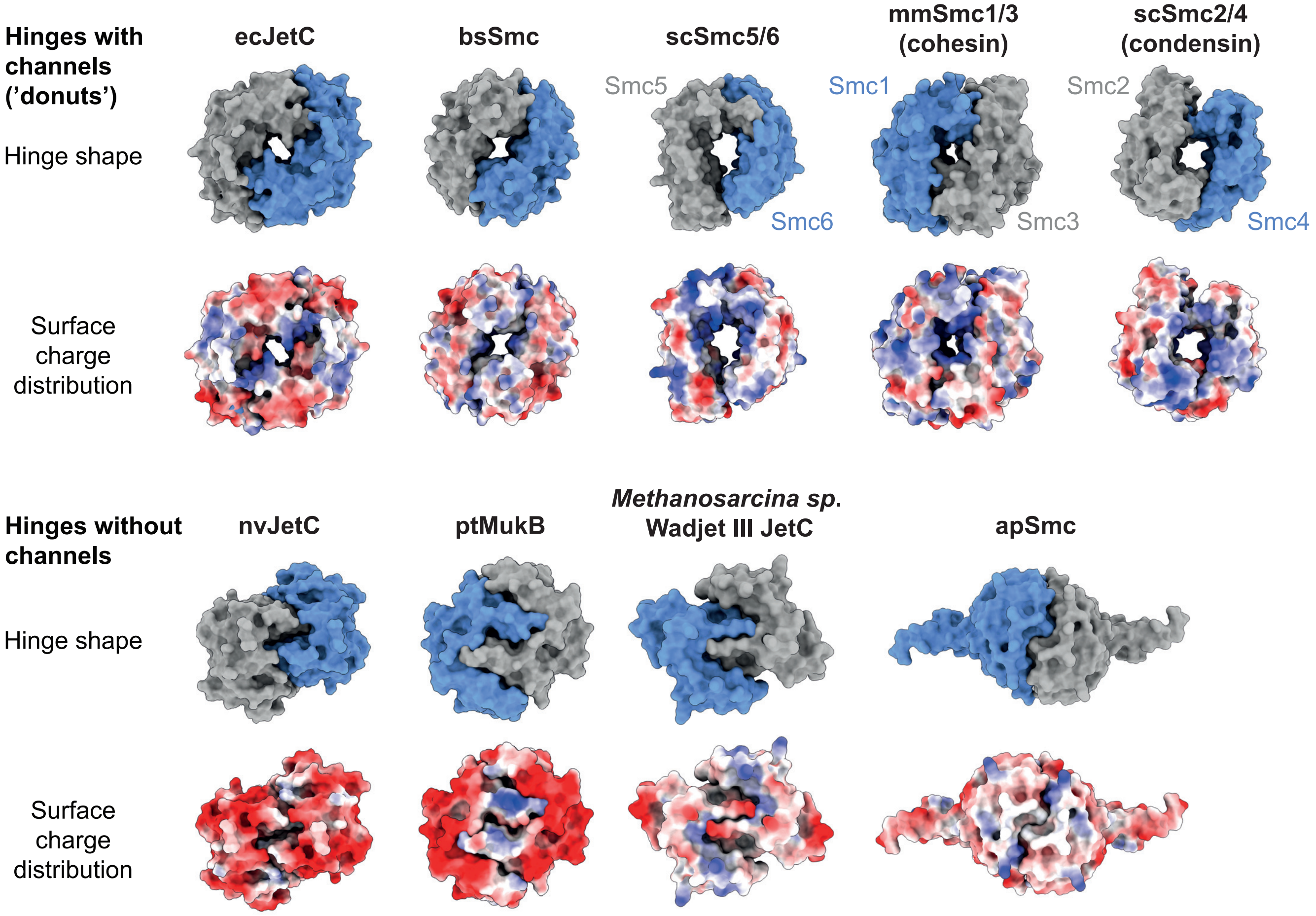


### Extended Data Fig. 8

Extended Data Fig. 8 - Hinge structures of selected SMC complexes.

Top: top view structures, bottom: surface charge distribution. Shown here are eight diverse selected SMC hinges, five “donut hinges” containing discernible channels and four without. The ecJetC, nvJetC, bsSmc, scSmc5/6, mmSmc1/3 and scSmc2/4 hinges are as shown in Fig. 3A. ScSmc2/4: *Saccharomyces cerevisiae* condensin (PDB:6YVU); mmSmc1/3: *Mus musculus* cohesin (PDB: 2WD5); ptMukB: *Photorhabdus thracensis* MukB (PDB: 7NYY); Wadjet-III JetC hinge from *Methanosarcina sp.* (AF3 prediction, NCBI accession number KKG07771.1); apSmc: *Acetobacter pasteurianus* Smc (AF3 prediction, NCBI accession number BAI20494.1). For the channel-less hinges of Wadjet-II and -III, the apparent loss of hinge bypass may have resulted from a Wadjet-specific molecular conflict, as donut-shaped hinges remain predominant outside the Wadjet-II and -III families. Accordingly, we also note that channel-less hinges tend to exhibit greater structural diversity, suggesting that hinges undergo accelerated evolutionary changes when the constraints imposed by hinge bypass are relaxed. Nevertheless, in the donut-shaped hinges, the gate dynamics might be fine-tuned to suit different SMC functions. Wadjet may prevent ssDNA (and ssDNA mimics) from stably adhering to the hinge to avoid interference with plasmid restriction, while other complexes may remain associated with ssDNA via the hinge for extended periods to facilitate DNA repair, as proposed for Smc5/6.
